# Cation-exchange synthesized Zn-doped Ag_2_S Nanostructures for Photothermal and Photodynamic Therapies across Breast Cancer Subtypes

**DOI:** 10.1101/2025.10.26.684667

**Authors:** Harshavardhan Mohan, Satabdi Acharya, Inhee Chung, Taeho Shin

## Abstract

Photothermal therapy (PTT) and photodynamic therapy (PDT) require nanostructures capable of efficiently converting red-light energy into both heat and reactive oxygen species (ROS). However, simultaneously achieving high photothermal conversion efficiency and strong ROS generation remains challenging. Here, we report zinc-doped Ag_2_S (ZSS) nanostructures, synthesized via controlled cation exchange, in which Zn incorporation modulates the electronic structure of Ag_2_S and improves charge separation, thereby enhancing red-light-activated PTT/PDT performance while preserving intrinsic biocompatibility. Among the series, ZSS(0.15) demonstrated optimized charge-carrier dynamics, a high photothermal conversion efficiency of 67.26%, and approximately four-fold higher singlet oxygen (^1^O_2_) generation relative to methylene blue under 660 nm irradiation. These physicochemical enhancements translated into potent therapeutic outcomes: *in-vitro*, ZSS(0.15) achieved an IC_50_ of 15 µg/mL under irradiation, corresponding to approximately a 1.5-fold enhancement in cytotoxic potency compared to pristine Ag_2_S, and induced apoptosis via activation of the p53/Bax/Caspase pathway. *In-vivo*, ZSS(0.15) with laser irradiation achieved ∼97% tumor volume suppression without systemic toxicity. Extending evaluation across multiple breast cancer cell lines representing distinct molecular subtypes further confirmed broad-spectrum *in-vitro* efficacy. Altogether, these findings demonstrate that controlled Zn incorporation into Ag_2_S effectively enhances red-light-driven photothermal conversion and ^1^O_2_ generation, establishing a rational materials strategy for improving synergistic PTT/PDT performance.

**GRAPHICAL ABSTRACT:** A Zn-doped Ag_2_S nanoplatform (ZSS(0.15)) enables synergistic photothermal and photodynamic therapy under 660-nm laser irradiation, triggering p53-mediated mitochondrial apoptosis via ROS generation and caspase-3 activation, achieving ∼97% tumor regression *in-vivo* without detectable systemic toxicity.

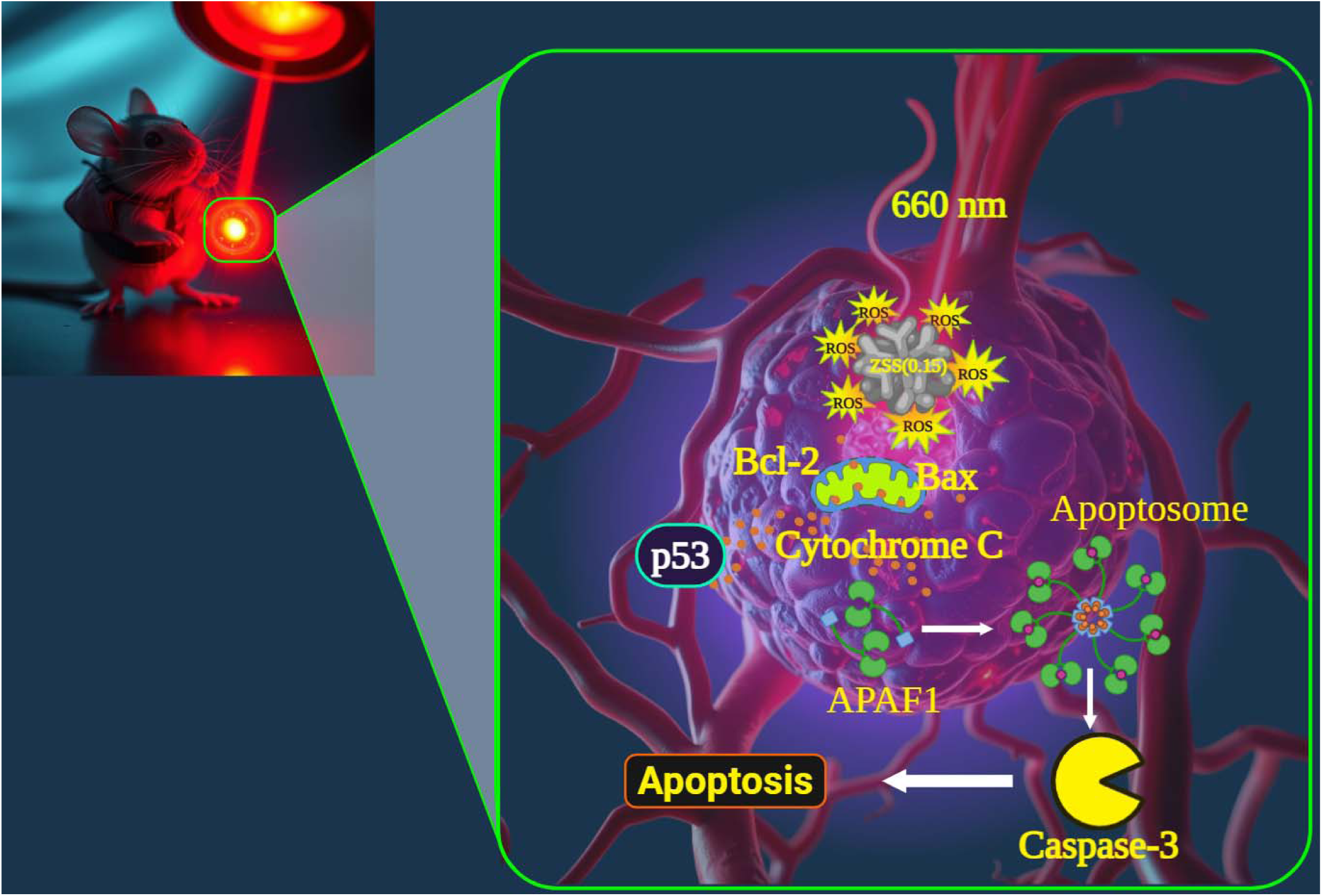

## 1. INTRODUCTION

Breast cancer remains one of the leading causes of cancer-related mortality in women worldwide [1]. Despite advances in chemotherapy, radiotherapy, and targeted molecular interventions, current treatments are constrained by systemic toxicity, limited tumor specificity, and the emergence of multidrug resistance [2]. The non-specific biodistribution of therapeutic agents and the accumulation of off-target toxicities not only reduce treatment efficacy but also compromise patient quality of life [3]. While conventional approaches can achieve disease control in certain cases, they frequently fail to fully eradicate tumors, thereby increasing the risk of recurrence [4, 5]. These limitations highlight the urgent need for alternative, minimally invasive treatment modalities that combine tumor-specific targeting with reduced collateral damage to healthy tissues.

Light-based therapies have emerged as attractive solutions to these challenges, offering spatial and temporal control of treatment with minimal side effects [6]. In particular, photothermal therapy (PTT) and photodynamic therapy (PDT) are considered synergistic and highly promising non-invasive approaches with reduced systemic toxicity [7]. PTT induces localized hyperthermia via light-activated photothermal agents, whereas PDT relies on photosensitizers to generate reactive oxygen species (ROS), including singlet oxygen (^1^O_2_), superoxide anions (O_2_^*-^), and hydroxyl radicals (*OH), which disrupt cellular redox homeostasis and trigger apoptotic cascades [8, 9]. Recent progress in nanotechnology has enabled the design of multifunctional nanomaterials that integrate both PTT and PDT into a single therapeutic platform [10, 11]. Among these, silver sulfide (Ag_2_S) nanostructures are particularly promising due to their strong red-light absorption, high photostability, and relatively low toxicity [12–14]. For instance, Liu et al. [15] demonstrated a tumor-inhibiting Ag_2_S based nanoplatform with substantial tumor reduction, but the system still required improvements in synthesis simplicity and overall therapeutic efficacy.

One effective strategy to overcome these shortcomings is metal-ion doping, which modulates the optical, electronic, and photothermal properties of Ag_2_S. Although various dopants have been investigated, many raise concerns regarding biosafety and long-term *in vivo* compatibility [16]. Zinc presents a particularly attractive alternative due to its essential biological role, low cytotoxicity at therapeutic doses, and documented metabolic clearance pathways. [17]. Moreover, Zn^2+^ incorporation into semiconductor lattices is known to induce lattice distortions and modulate the electronic structure (including defect states), thereby promoting efficient charge separation and suppressing radiative recombination key factors for improving both light-to-heat conversion and ROS generation [18]. Despite these advantages, the therapeutic potential of Zn-doped Ag_2_S nanostructures has not been systematically investigated.

In this study, we synthesized and characterized Zn-doped Ag_2_S (ZSS) nanostructures with systematically varied doping levels (Zn:Ag, 0.0 to 0.30) to identify the optimal composition that balances structural integrity, charge separation, and therapeutic efficacy under 660 nm red-light irradiation. The objectives were to elucidate the physicochemical and photo-responsive behavior of ZSS nanostructures, evaluate their ROS-mediated cytotoxicity, and investigate the underlying mechanistic pathways *in-vitro* and *in-vivo*. To establish broad spectrum relevance, the therapeutic potential was assessed across multiple breast cancer cell lines representing distinct molecular subtypes [19, 20]: luminal A (MCF-7, T47D), HER2-positive (SK-BR-3), and triple-negative (MDA-MB-231). Altogether, this study evaluates Zn incorporation as a rational strategy to enhance Ag_2_S based red-light-activated synergistic PTT/PDT, providing mechanistic and therapeutic insight into dopant-mediated optimization of semiconductor nanostructures for cancer phototherapy.

## 2. RESULTS AND DISCUSSION

### 2.1. Structural and compositional characterization

#### 2.1.1. XRD

The crystallographic features of the synthesized Zn_x_Ag_2-2x_S (ZSS) hybrid nanostructures (**Figure 1a**) with varying Zn-to-Ag ratios (x = 0.05-0.30) were investigated using X-ray diffraction (XRD, MiniFlex 600, Korea I.T.S Co., Ltd), as shown in **Figure 1b**. The pristine Ag_2_S sample exhibited characteristic orthorhombic acanthite phase peaks indexed to (101), (−112), (−121), (120), (112), (031), and (200) planes (JCPDS No. 14-0072), confirming phase purity [21]. Conversely, the ZnS sample displayed a cubic zinc blende phase with diffraction peaks corresponding to (111), (200), (220), and (311) planes (JCPDS No. 05-0566), consistent with its well-known crystal structure [22]. Upon progressive substitution of Zn^2+^ into the Ag_2_S lattice (ZSS(0.05) - ZSS(0.30)), all samples up to ZSS(0.25) retained the orthorhombic Ag_2_S phase with no secondary phases detected, suggesting successful solid-state incorporation of Zn^2+^ ions without significant lattice distortion. However, at a Zn:Ag ratio of 0.30:1.70 (ZSS(0.30)), additional peaks matching the ZnS phase emerged, indicating phase segregation and a solubility threshold beyond which excess Zn precipitates out as a secondary ZnS phase. Therefore, ZSS(0.25) was identified as the composition exhibiting the highest Zn incorporation while maintaining single-phase structural integrity, enabling effective compositional tuning without phase separation. The phase-pure nature of ZSS is critical for maintaining homogeneous energy band structures and interfacial charge transfer pathways, which directly influence its photophysical and therapeutic properties, as discussed in subsequent sections. The absence of secondary phases in ZSS(0.05) - ZSS(0.25) suggests a well-integrated substitutional doping mechanism, where Zn^2+^ ions, owing to their smaller ionic radius (0.74 Å) compared to Ag^+^ (1.15 Å), substitute Ag^+^ sites without significantly distorting the host lattice [22]. Notably, the gradual shift and slight broadening of diffraction peaks with increasing Zn content (especially visible between ZSS(0.10) - ZSS(0.25)) indicate minor lattice compression, affirming successful Zn incorporation and nanoscale crystallinity. These findings validate the rational design strategy adopted in this study, where phase purity was prioritized to ensure predictable and improved photothermal and photodynamic behavior.

**Figure 1.**
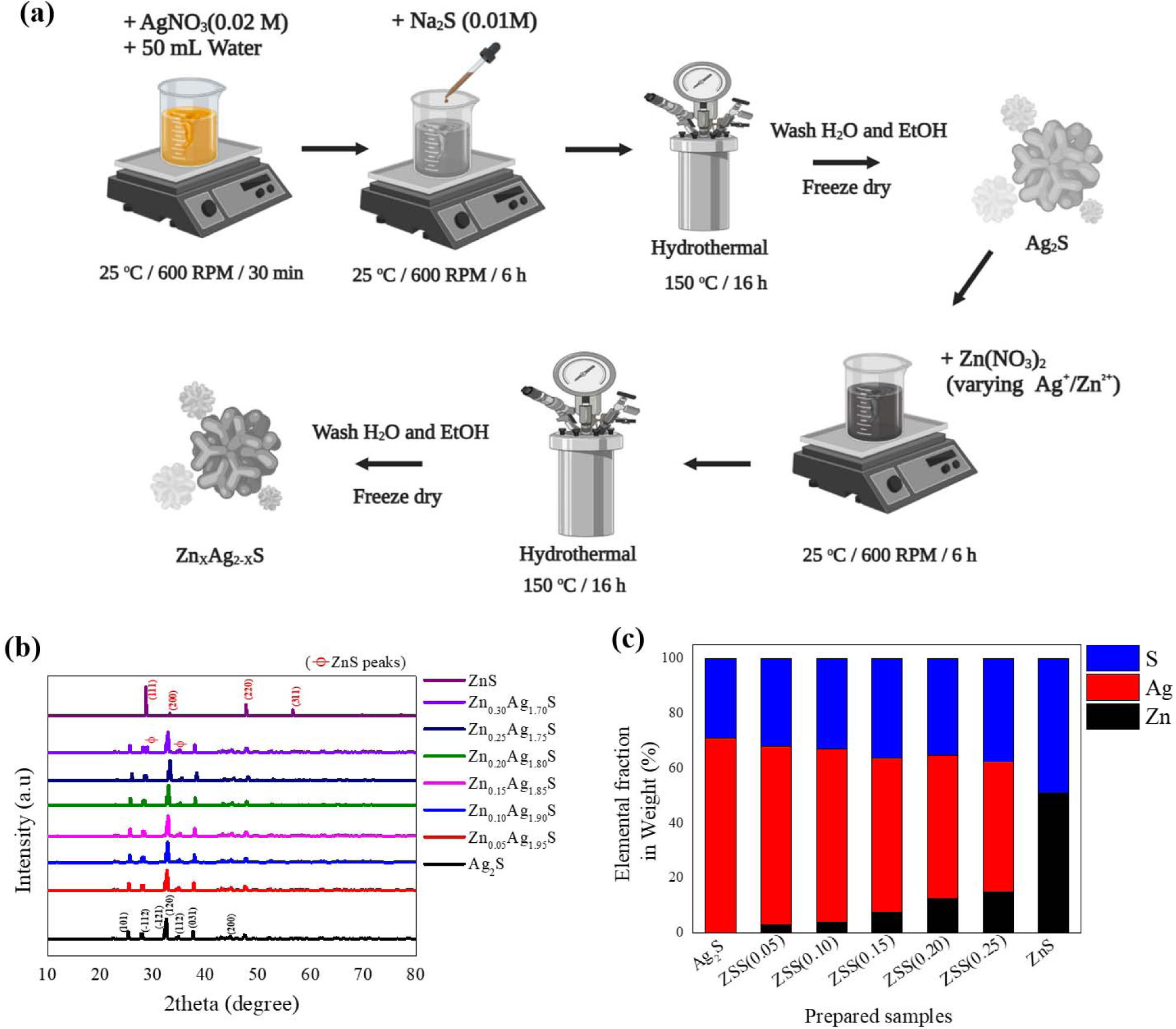
**(a)** Schematic illustration of the synthesis procedure for Zn_x_Ag_2-2x_S (ZSS) nanostructures; **(b)** XRD patterns of the synthesized ZSS samples; **(c)** Elemental composition (wt%) obtained from ICP-OES analysis.

#### 2.1.2. ICP-OES

To verify the successful incorporation of Zn into the Ag_2_S lattice and to quantify the elemental distribution, inductively coupled plasma optical emission spectroscopy (ICP-OES (Agilent 5800)) was performed for all synthesized samples, as shown in **Figure 1c**. The pristine Ag_2_S sample exhibited a binary composition of Ag and S with no detectable Zn content, consistent with its nominal stoichiometry. With the gradual incorporation of Zn (ZSS(0.05) - ZSS(0.25)), a clear trend of increasing Zn weight percentage was observed, accompanied by a proportional decrease in Ag content. This confirms the intended cation exchange between Ag and Zn ions, supporting the substitutional doping strategy. For instance, ZSS(0.05) and ZSS(0.10) showed minor Zn incorporation (<5 wt%), reflecting a low but successful doping level. As the Zn ratio increased, samples ZSS(0.15) and ZSS(0.20) demonstrated moderate Zn contents (∼10-15 wt%) with a corresponding reduction in Ag, suggesting a balanced doping regime. Notably, ZSS(0.25) exhibited the highest Zn incorporation (∼20 wt%) while retaining significant Ag, consistent with its XRD pattern indicating a single-phase solid solution without phase segregation. In contrast, the ZnS reference showed a complete absence of Ag, confirming its phase purity. The progressive elemental shift from Ag-rich to Zn-rich compositions across the ZSS series aligns well with the designed stoichiometric ratios. These quantitative findings complement the structural insights from XRD, further validating the formation of compositionally tunable Zn_x_Ag_2-2x_S hybrids.

#### 2.1.3. TEM-EDS

To gain insight into the structural features and elemental distribution of the ZSS hybrids, transmission electron microscopy (TEM, Hitachi JEOL-2010 HR-TEM.) and energy-dispersive X-ray spectroscopy (EDS) analyses were performed for ZSS(0.15) as presented in **Figure 2**. Low-magnification TEM images (**Figure 2a**-**b**) reveal a crumpled, petal-like morphology indicative of ultrathin, wrinkled nanosheets aggregated into hierarchical flower-like structures. High-resolution TEM (**Figure 2c**) highlights the crystalline lattice fringes with an interplanar spacing of 0.330 nm, corresponding to the (101) plane of the Ag_2_S crystal structure. These results support the well-defined crystallinity and anisotropic growth of the ZSS(0.15) nanostructures. The selected area electron diffraction (SAED) pattern (**Figure 2d**) exhibits multiple concentric rings indexed to the (101), (112), (120), (103), and (220) planes, confirming the polycrystalline nature of the material, complementing XRD data (**Figure 1b**). The bright-field scanning TEM image (**Figure 2e**) further supports the 3D interconnected nanosheet architecture. Elemental mapping through EDS (**Figures 2f-h**) clearly shows the homogeneous distribution of Zn (red), Ag (cyan), and S (green) throughout the structure, signifying successful doping and uniform incorporation of Zn into the Ag_2_S matrix without phase segregation. The EDS quantification table confirms an atomic composition of Zn: 7.49%, Ag: 56.26%, and S: 36.25%, consistent with the intended stoichiometry and the ICP-OES data (**Figure 1c**). TEM-derived size statistics (n = 75), as shown in **Figure SF1**, exhibit a Gaussian distribution with a dominant peak centered around 160 nm, indicating a relatively narrow size distribution. Most of the nanostructures fall within the 140-180 nm range, confirming the formation of uniformly distributed ZSS nanoflowers. This size range is ideal for biomedical applications, especially for enhanced tumor accumulation via the enhanced permeability and retention effect, while still small enough to promote efficient cellular uptake [23].

**Figure 2.**
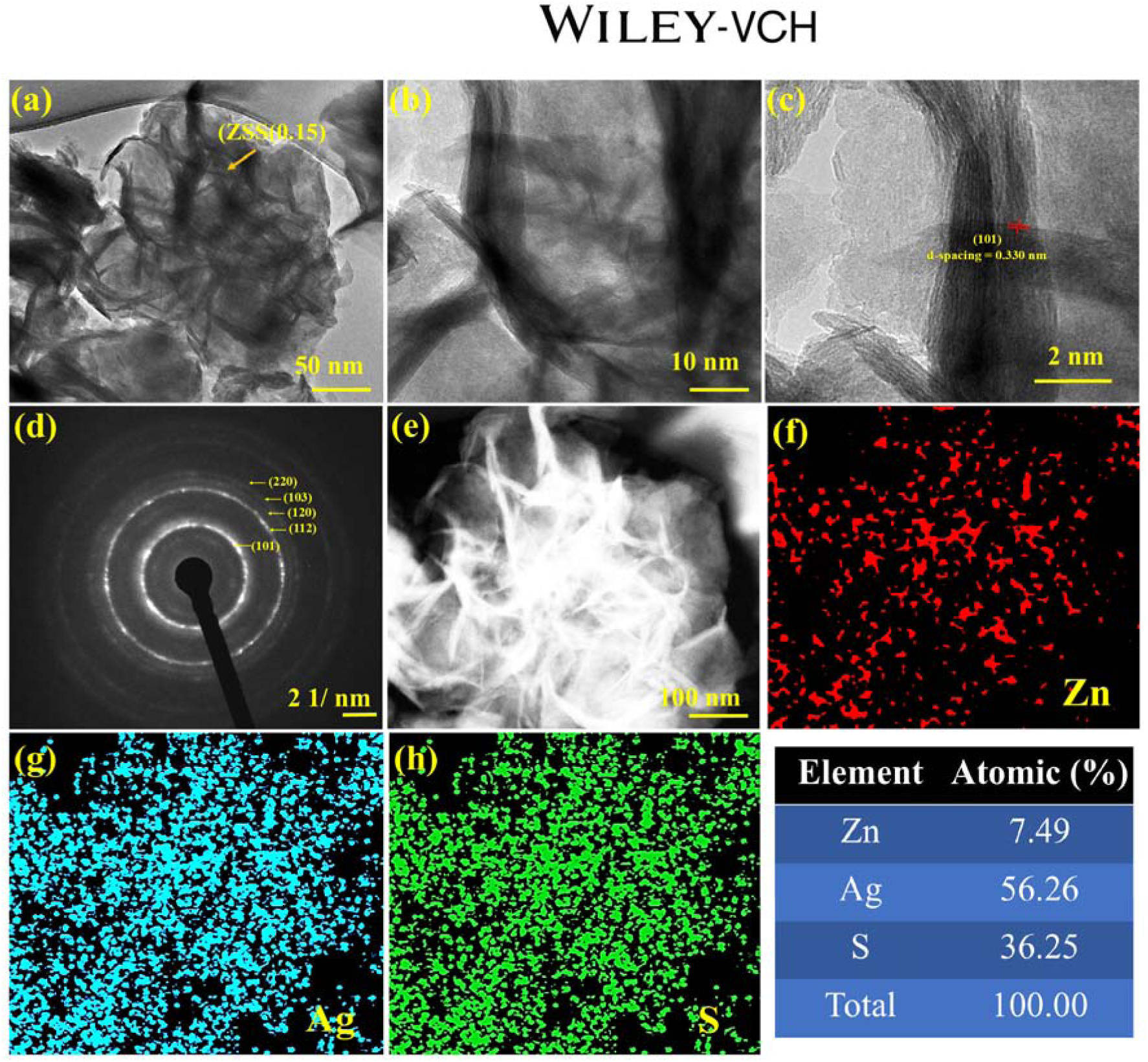
**(a, b)** Low-magnification TEM images of ZSS(0.15) nanostructures; **(c)** High-resolution TEM (HRTEM) image; **(d)** Corresponding SAED pattern; **(e)** Bright-field STEM image; **(f-h)** Elemental mapping of Zn, Ag, and S obtained from EDS analysis.

### 2.2. Optical and Electronic Properties

#### 2.2.1. DRS-UV

The optical properties of ZSS nanostructures were systematically investigated using UV-Vis diffuse reflectance spectroscopy (DRS-UV-NIR, Jasco V-770 UV-Vis-NIR), and the corresponding Tauc plot (**Figure 3a**) revealed a clear shift in optical absorption with increasing Zn content. As the Zn:Ag ratio increased from 0.00 to 0.25, the optical bandgap widened progressively from 2.04LeV (pure Ag_2_S) to 2.50LeV, approaching the wider bandgap of ZnS (3.27LeV). This blue shift in the absorption edge arises from the incorporation of smaller Zn^2+^ ions into the Ag_2_S lattice, inducing lattice strain and electronic structure perturbations that change the electronic structure [24, 25]. This absorption behavior suggests that sub-bandgap defect-state transitions may contribute to absorption in the red region (660 nm) [26]. Zn incorporation can affect the optical absorption and charge carrier dynamics of the nanostructures, therefore potentially contributing to enhanced ROS generation and photothermal performance under 660 nm irradiation. This composition-dependent optical response highlights the importance of controlled Zn incorporation in optimizing red-light responsiveness for synergistic PDT/PTT applications.

**Figure 3.**
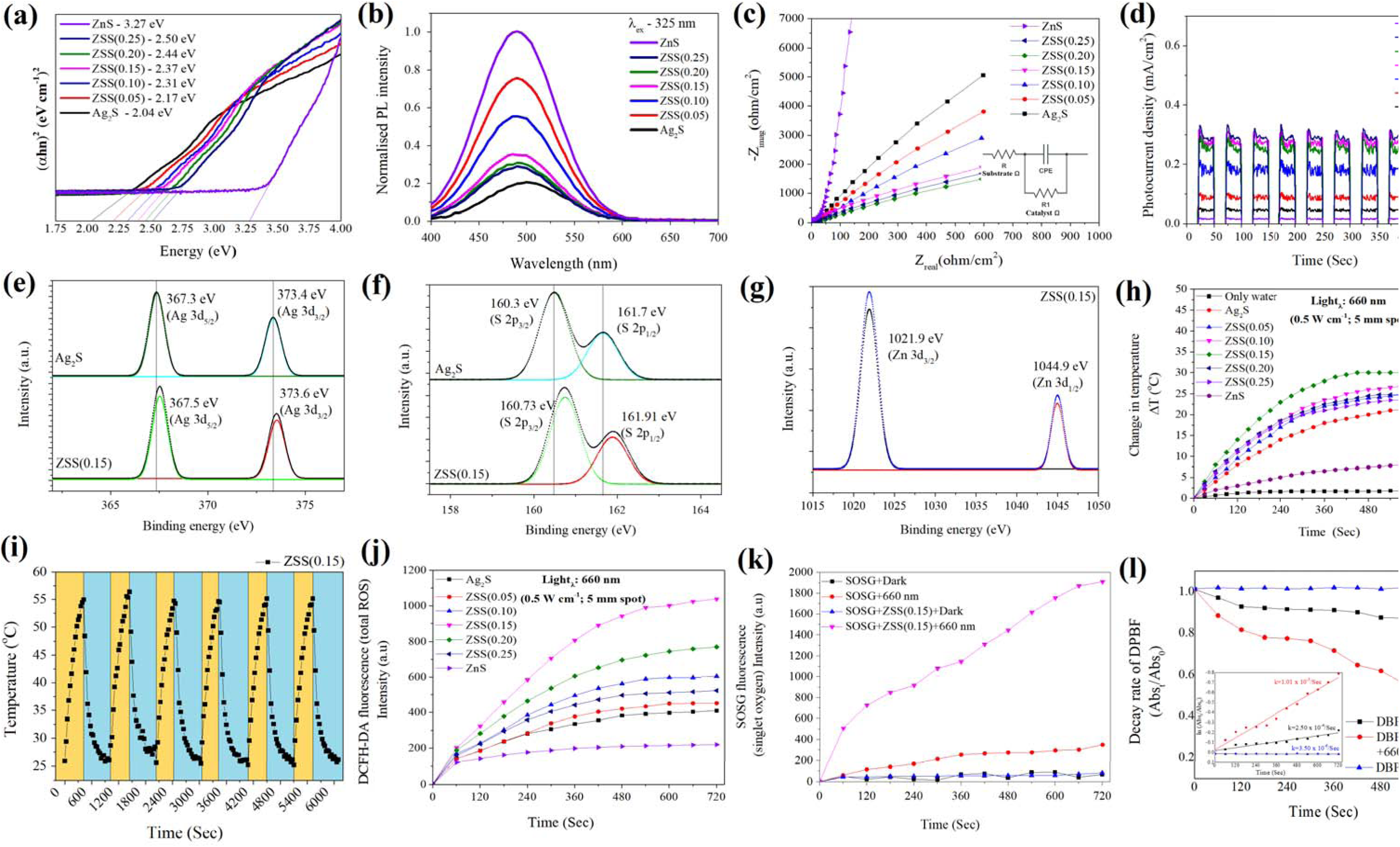
**(a)** UV-Vis diffuse reflectance spectra (DRS); **(b)** Photoluminescence (PL) spectra; **(c)** Electrochemical impedance spectroscopy (EIS) plots; **(d)** Transient photocurrent response; **(e-g)** High-resolution XPS spectra of Ag 3d, S 2p, and Zn 2p, respectively; Photothermal **(h)** heating curves and **(i)** stability; **(j)** Time-dependent ROS generation profiles of the prepared nanostructures; **(k)** Singlet oxygen (^1^O_2_) generation of ZSS(0.15) in light and dark condition; **(l)** DPBF photobleaching efficiency and kinetics **(inset)** of ZSS(0.15) with MB as reference photosensitizer

#### 2.2.2. Photoluminescence

The photoluminescence (PL, Horiba Fluoromax-4 spectrofluorometer) spectra of the ZSS nanostructures (**Figure 3b**), excited at 325 nm, reveal distinct emission intensity variations with progressive Zn incorporation, reflecting the evolution of charge carrier dynamics. Pure ZnS exhibits the highest PL intensity, indicative of high radiative recombination rates, whereas pure Ag_2_S shows the lowest intensity, suggesting efficient charge separation. As Zn doping increases, a non-linear trend emerges, with initial doping (e.g., ZSS(0.05)) leading to increased PL due to defect-related trap states, while higher doping levels (e.g., ZSS(0.15) - ZSS(0.25)) result in PL quenching. This reduction in emission intensity at moderate-to-high Zn content is consistent with suppressed electron/hole recombination (potentially with increased non-radiative relaxation), likely owing to enhanced charge separation and defect-state passivation [27]. Such behavior is highly desirable for both photodynamic and photothermal therapy, as lower recombination rates prolong the lifetime of photogenerated charge carriers, thereby facilitating more efficient ROS generation and thermal energy conversion [28]. It should be noted that the PL analysis herein is intended to qualitatively evaluate charge recombination/separation behavior, rather than to quantify fluorescence quantum yield or imaging performance. These findings confirm that rational band engineering via Zn substitution can strategically manipulate carrier dynamics, optimizing the nanostructures for dual-modal PDT/PTT cancer therapeutic applications.

#### 2.2.3. EIS

Electrochemical impedance spectroscopy (EIS, ZIVE SP1, WonATech, Korea, with a standard three-electrode system) analysis was employed to investigate the interfacial charge transfer characteristics of the ZSS nanostructures (**Figure 3c**), with the Nyquist plots clearly highlighting significant differences in charge transfer resistance (R_ct_) among the samples. Pure ZnS exhibited the largest semicircle, corresponding to the highest R_ct_, which reflects its inherently poor conductivity and sluggish interfacial charge transfer. Conversely, Ag_2_S displayed markedly lower resistance, indicative of improved electrical conductivity [29]. Upon Zn incorporation, a progressive decline in R_ct_ was observed up to a certain doping level (**Table ST3**), signifying enhanced electron mobility and interfacial conductivity. This trend is particularly evident for intermediate ZSS compositions (e.g., ZSS(0.15) - (0.20)), where minimized charge transfer resistance suggests a synergistic balance between defect engineering and lattice conductivity. These findings are consistent with the PL spectra, which show reduced photoluminescence intensity for the same compositions, indicating suppressed radiative recombination and more efficient separation of charge carriers, which contributes to enhanced photo-induced reactive oxygen species generation rather than enhanced fluorescence emission.

#### 2.2.4. Photocurrent

The transient photocurrent measurements (**Figure 3d**, ZIVE SP1, WonATech, Korea, with a standard three-electrode system) further substantiate the charge separation and transport dynamics of the ZSS nanostructures under light irradiation. Upon periodic light on/off cycles (SOLIS-1C, Thorlabs, USA), a sharp rise and decay in photocurrent density were observed, with intensity varying across the compositional series. Pure ZnS showed minimal response (∼0.05 μA/cm^2^), attributed to its wide bandgap and inefficient charge separation, while Ag_2_S exhibited a moderate signal (∼0.18 μA/cm^2^) due to its narrower bandgap yet higher recombination. Remarkably, the ZSS nanoflowers, especially ZSS(0.15) and ZSS(0.20), displayed significantly enhanced photocurrent responses of ∼0.38 μA/cm^2^ and ∼0.35 μA/cm^2^, respectively, indicating superior photoinduced charge generation and extended carrier lifetimes. These trends are well-supported by EIS measurements, where the charge transfer resistance (R_ct_) for ZSS(0.15) and ZSS(0.20) was the lowest (∼820 and ∼860 Ω·cm^2^, respectively), in contrast to ZnS (∼6800 Ω·cm^2^) and Ag_2_S (∼5100 Ω·cm^2^), reflecting more efficient interfacial charge transport. Concurrently, these samples also exhibited suppressed PL intensity, compared to ZnS and Ag_2_S, confirming reduced radiative recombination. Collectively, the synergy between high photocurrent response, low R_ct_, and quenched PL intensity highlights the optimized charge carrier behavior achieved through optimal Zn substitution, ultimately promoting efficient ROS generation and photothermal conversion, both critical to enhancing therapeutic performance in PDT and PTT.

#### 2.2.5. XPS

To investigate the electronic structure of the prepared materials, high-resolution XPS spectra (Omicron XPS, Scienta Omicron, Germany) were acquired for pristine Ag_2_S and ZSS(0.15) nanostructures. The Ag 3d region of Ag_2_S (**Figure 3e**) reveals two strong peaks located at 367.3 eV and 373.4 eV, assigned to Ag 3d_5/2_ and Ag 3d_3/2_, respectively. These binding energies are consistent with Ag^+^ in Ag_2_S [30]. Upon incorporation of Zn into the lattice, the Ag 3d peaks in ZSS(0.15) exhibit a slight positive shift to 367.5 eV and 373.6 eV. This upward shift in binding energy indicates a slight increase in the effective nuclear charge experienced by the Ag atoms, which can be attributed to electronic interactions and lattice distortion induced by Zn substitution. The shift reflects a partial withdrawal of electron density from the Ag atoms, likely resulting from charge redistribution between Zn^2+^ and Ag^+^ species in the lattice. The S 2p core-level spectra (**Figure 3f**) further supports this electronic modulation. In Ag_2_S, the doublet peaks appear at 160.3 eV (S 2p_3/2_) and 161.7 eV (S 2p_1/2_), corresponding to divalent sulfide (S^2-^) ions. For ZSS(0.15), the S 2p peaks are shifted slightly to higher binding energies, located at 160.73 eV and 161.91 eV, respectively. This subtle shift implies that the sulfur atoms experience a modified chemical environment due to neighboring Zn dopants, which may cause local lattice strain and altered electronic density around sulfur atoms. Such a shift also supports the hypothesis of a strong interaction between Zn and the Ag_2_S matrix, rather than mere physical mixing. The Zn 2p spectrum (**Figure 3g**) of ZSS(0.15) displays well-defined peaks at 1021.9 eV (Zn 2p_3/2_) and 1044.9 eV (Zn 2p_1/2_), characteristic of Zn^2+^ species in a sulfide matrix [31]. The absence of any satellite peaks rules out the presence of metallic Zn or ZnO impurities, confirming that Zn is successfully incorporated into the Ag_2_S framework via cationic substitution rather than surface adsorption. This substitution is further validated by the consistent energy separation (∼23 eV) between Zn 2p_3/2_ and Zn 2p_1/2_ peaks, corresponding to characteristic spin-orbit splitting of divalent Zn^2+^ [32].

### 2.3. Photothermal and Photodynamic Performance

#### 2.3.1. Photothermal activity

To evaluate the photothermal response of the synthesized ZSS nanostructures, temperature elevation studies were performed under continuous 660 nm laser irradiation (0.5 W/cm^2^, 5 mm spot; Techhood (660 nm) word-line laser module, 300 mW max power, HK). The temperature change (ΔT) of various samples, including pristine Ag_2_S and ZSS nanostructures was monitored over a 12-minute irradiation period. As shown in **Figure 3h**, all materials exhibited a gradual increase in temperature (Ebro TFI 54 Infrared Thermometer) over time, but with significantly varying efficiencies depending on the Zn substitution level. Among all tested compositions, the ZSS(0.15) sample demonstrated the most pronounced photothermal response, with a maximum ΔT of 30 °C, whereas the pristine Ag_2_S exhibited only a ΔT of ∼22 °C under identical conditions. The enhancement in photothermal performance upon Zn incorporation can be attributed to synergistic improvements in light absorption and non-radiative relaxation pathways introduced by Zn-mediated band structure modulation [33]. To gain further insights, heating and cooling profiles of ZSS(0.15) were recorded (**Figure SF2**), wherein the sample temperature rose from ambient (∼25 °C) to a plateau of 55 °C within 12 minutes and returned close to the baseline after stopping irradiation. From the cooling profile, the photothermal conversion efficiency (η) was calculated to be 67.26%, indicating excellent light-to-heat conversion properties. For photothermal stability assessment, ZSS(0.15) dispersions were subjected to repeated laser ON/OFF cycling. Briefly, the laser was switched ON until the temperature reached a steady maximum, followed by switching OFF until the solution returned to baseline temperature.

This process was repeated for 6 consecutive cycles under identical irradiation conditions (**Figure 3i**). Notably, no obvious decay in maximum temperature elevation or heating rate was observed across successive cycles, demonstrating excellent photothermal stability and reproducibility under repeated laser exposure. Collectively, these observations suggest that Zn incorporation at 0.15 induces optimal defect states and band alignment, which not only improve charge separation and transport but also enhance non-radiative recombination routes, crucial for photothermal conversion [34]. The improved light-harvesting ability and reduced electron-hole recombination rates likely contribute to the higher photothermal output of ZSS(0.15), making it the most efficient composition for further photodynamic investigations.

#### 2.3.2. ROS generation and singlet oxygen (^1^O_2_) evaluation under 660 nm irradiation

The generation of photo-induced ROS under 660 nm laser irradiation (0.5 W cm^-2^, 5 mm spot) was first evaluated using the DCFH-DA fluorescence assay as a general indicator of oxidative stress (**Figure 3j**). The fluorescence intensity increased progressively with irradiation time for all tested samples, confirming light-triggered ROS production. Notably, ZSS(0.15) exhibited the highest fluorescence intensity at all time intervals, reaching ∼1038 a.u. at 720 s, significantly surpassing Ag_2_S, ZnS, and other Zn-doped compositions. The ROS generation trend followed the order: ZSS(0.15) > ZSS(0.20) > ZSS(0.10) > ZSS(0.25) > ZSS(0.05) > Ag_2_S > ZnS. It should be noted that DCFH-DA is not specific to a particular ROS type and therefore reflects overall oxidative activity rather than singlet oxygen generation. To identify the dominant ROS species involved in the photodynamic process, singlet oxygen (^1^O_2_) generation was subsequently examined using Singlet Oxygen Sensor Green (SOSG) (**Figure 3k**). Under 660 nm irradiation, ZSS(0.15) induced a pronounced increase in SOSG fluorescence intensity, whereas negligible signal changes were observed under dark conditions or in control samples. This result directly confirms ^1^O_2_ as a primary ROS species generated by ZSS(0.15) upon red-light irradiation. Furthermore, the relative efficiency of ^1^O_2_ generation was quantitatively assessed using a DPBF photobleaching assay with methylene blue as a reference photosensitizer (**Figure 3l**). The temporal decay of DPBF absorbance followed pseudo-first-order kinetics (**Figure 3l-inset**), as evidenced by the linear relationship between ln(A_t_/A_0_) and irradiation time. Under identical irradiation conditions, ZSS(0.15) exhibited a significantly higher DPBF degradation rate constant (k = 1.01 × 10^-3^ s^-1^) compared to MB (k = 2.50 × 10^-4^ s^-1^), corresponding to an approximately four-fold enhancement in relative ^1^O_2_ generation efficiency. In contrast, DPBF alone showed negligible decay (k = 3.5 × 10^-6^ s^-1^), confirming that photobleaching was dominated by photo-induced ^1^O_2_ rather than self-degradation.

These findings correlate well with the photocurrent, EIS, and PL results, where ZSS(0.15) demonstrated superior charge separation efficiency, reduced interfacial resistance, and suppressed radiative recombination. Collectively, the integration of general ROS assessment (DCFH-DA), ^1^O_2_-specific identification (SOSG), and relative ^1^O_2_ quantification (DPBF) confirms that ZSS(0.15) achieves an optimized photodynamic response while maintaining strong photothermal performance, thereby offering a balanced and enhanced dual-mode therapeutic platform.

### 2.4. In-vitro study

#### 2.4.1. MTT-assay

To evaluate the phototherapeutic efficacy of the synthesized ZSS nanostructures, an MTT assay was conducted against MCF-7 breast cancer cells under both light irradiation (660 nm, 0.5 W cm^-2^, 5 mm spot, 10 min) and dark conditions, across a concentration range of 0-25 μg/mL. All samples were compared with clinically used anticancer drug cisplatin as a control. Upon laser exposure (**Figure 4a**), all ZSS samples exhibited a clear dose-dependent cytotoxic response. Notably, ZSS(0.15) consistently outperformed the other compositions, showing significantly enhanced cell-killing efficiency. At a concentration of 25 μg/mL, ZSS(0.15) reduced the cell viability to ∼12%, compared to 22% for ZSS(0.10), 30% for ZSS(0.05), and 38% for pristine Ag_2_S. ZnS, lacking significant visible light absorption or charge separation ability, maintained high viability (∼84%), serving as a negative reference. This enhanced activity is attributed to the synergistic photothermal (PTT) and photodynamic (PDT) mechanisms of ZSS nanostructures. The optimal Zn doping (0.15 molar ratio) likely promotes superior light absorption, efficient electron-hole separation, and intensified ROS generation consistent with the previously discussed results from photothermal measurements, photocurrent response, and ROS activity assay. In the absence of light (**Figure 4b**), all nanostructures displayed markedly lower cytotoxicity, with cell viability remaining above 75% even at the highest dose (25 μg/mL). For ZSS(0.15), cell viability was ∼78%, affirming its low dark toxicity and strong photo-responsiveness indicating that the therapeutic effect is primarily light-triggered, confirming the biocompatibility of the system in non-irradiated conditions. Cisplatin, a well-established chemotherapeutic agent, showed robust cytotoxicity with an IC_50_ of ∼10 μg/mL, reducing cell viability to ∼9.1% at 25 μg/mL (**Figure 4c**).

**Figure 4.**
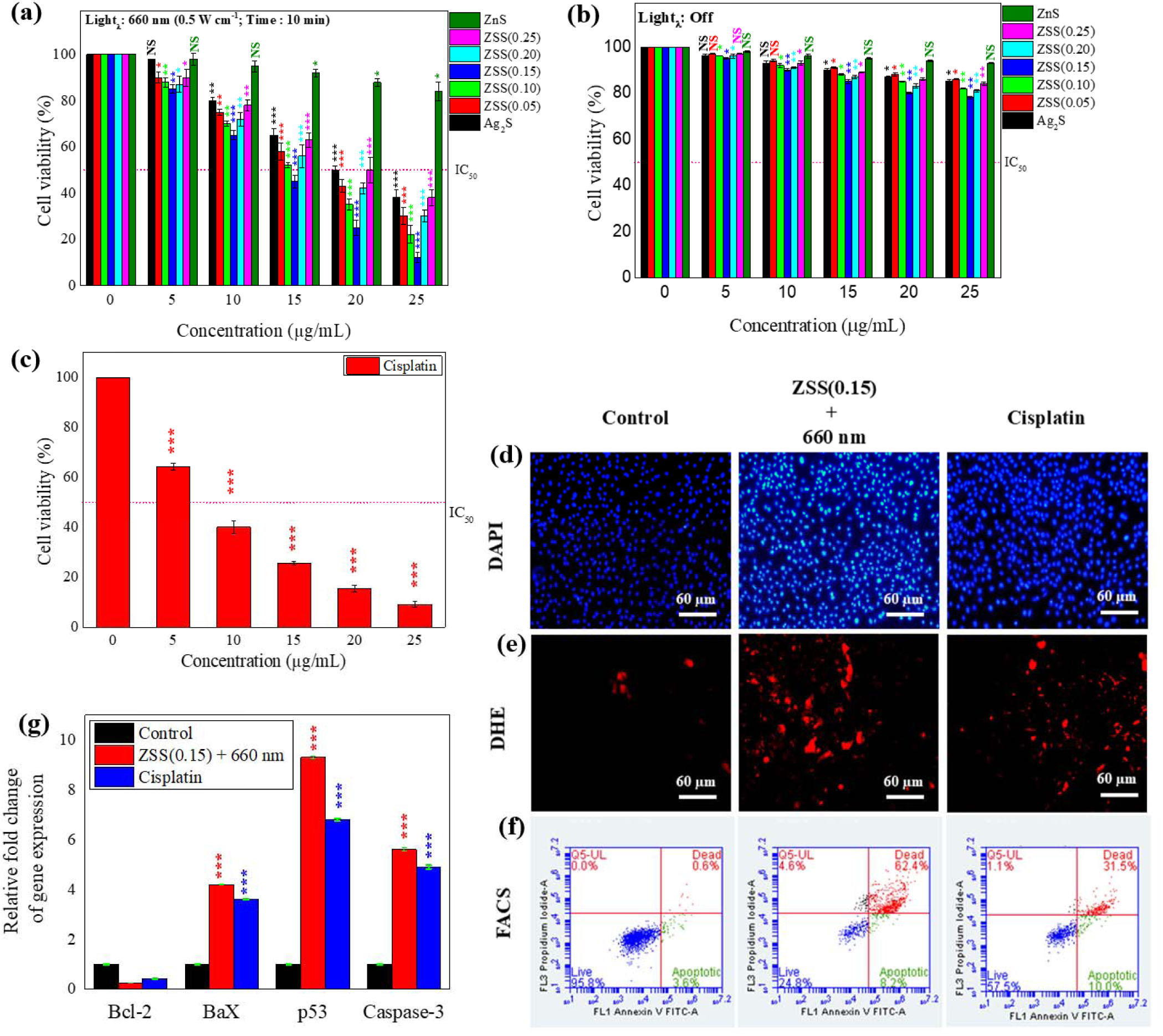
**(a)** Cell viability (MTT assay) under combined photothermal and photodynamic treatment (PTT/PDT); **(b)** Cell viability under dark conditions; **(c)** Cytotoxic effect of cisplatin; [Asterisks indicate statistically significant differences from 0 µg/mL within each treatment group; NS = no significance; * = *p* < 0.05; ** = *p* < 0.01; *** = *p* < 0.001] **(d)** DAPI staining for nuclear morphology; **(e)** DHE staining for intracellular ROS generation; **(f)** Flow cytometry (FACS) plots for apoptotic profiling; **(g)** Relative gene expression level [Asterisks indicate statistically significant differences from control within each group; NS = no significance; * = *p* < 0.05; ** = *p* < 0.01; *** = *p* < 0.001].

Interestingly, ZSS(0.15) closely matched this performance under laser irradiation, achieving an IC_50_ at ∼15 μg/mL, highlighting its potential as an effective alternative. Based on these findings, it can be concluded that the ZSS nanostructures demonstrate excellent photodynamic and photothermal therapy (PDT/PTT) capabilities, with ZSS(0.15) emerging as the most promising composition. Therefore, ZSS(0.15) at 15 μg/mL (IC_50_) was selected as the optimal concentration for all subsequent *in-vitro* biological experiments.

#### 2.4.2. Cell staining & FACS

To further validate the cytotoxic effects observed in the MTT assay, nuclear integrity and intracellular ROS generation were assessed through DAPI and DHE staining (detailed staining procedures are described in the *Supporting Information* file), respectively (**Figure 4d-f**), while apoptotic progression was quantified using flow cytometry (FACS). As shown in the DAPI images, all treatment groups exhibited uniform nuclear morphology; however, the ZSS(0.15)+660 nm and Cisplatin groups showed signs of nuclear condensation and shrinkage, indicative of apoptotic changes [35]. DHE staining revealed minimal ROS presence in the control group, while a dramatic increase in red fluorescence intensity was observed in ZSS(0.15)+660 nm treated cells, suggesting robust intracellular ROS generation upon light irradiation. This confirms the effective photodynamic functionality of the ZSS(0.15) nanostructures. The ROS levels in the Cisplatin group were comparatively lower than ZSS(0.15)+660 nm, further underscoring the superior oxidative stress induced by the photodynamic/photothermal therapy. Flow cytometry analysis provided quantitative insights into cell fate. The control group showed a predominance of live cells (95.8%) with negligible apoptosis (3.6%) and death (0.6%). In contrast, the ZSS(0.15)+660 nm group exhibited a significant increase in apoptotic (8.0%) and dead (62.4%) populations, with live cells reduced to 24.8%. Notably, this cell death rate was substantially higher than that of the Cisplatin group, which showed 31.5% dead and 10.1% apoptotic cells, indicating the enhanced therapeutic efficacy of the ZSS(0.15)+660 nm. Collectively, these *in-vitro* results confirm that ZSS(0.15) under 660 nm irradiation induces substantial ROS-mediated cytotoxicity and apoptosis, in line with the MTT findings [36]. The elevated ROS levels and apoptotic cell fractions support the proposed photodynamic therapeutic mechanism and warrant further exploration into associated apoptotic gene expression profiles [36].

#### 2.4.3. Gene expression

To gain deeper insights into the molecular mechanism underlying the observed anticancer efficacy of the nanostructured system, the expression profiles of key apoptosis-related genes BCL2, BAX, P53, and CASP3, were evaluated (detailed experimental conditions are described in the *Supporting Information* file) in MCF-7 cells following treatment with ZSS(0.15) under 660 nm irradiation (15 μg/mL) and compared with cisplatin (10 μg/mL), both at their respective ICLL concentrations. As shown in **Figure 4g**, treatment with ZSS(0.15)+laser resulted in a marked downregulation of BCL2 (0.25-fold), a key anti-apoptotic gene [37, 38]. This reduction was significantly greater than that observed with cisplatin (0.42-fold), indicating a stronger inhibition of pro-survival signaling. Simultaneously, the pro-apoptotic genes were dramatically upregulated in response to ZSS(0.15). Specifically, BAX increased to 4.2-fold, P53 to 9.3-fold, and CASP3 to 5.6-fold, compared to respective fold increases of 3.6, 6.8, and 4.9 in cisplatin-treated cells. These data indicate that ZSS(0.15) activated the intrinsic apoptotic pathway more effectively than cisplatin, with a strong emphasis on mitochondrial dysfunction and P53-mediated transcriptional activity. The robust upregulation of P53, a central regulator of cellular stress responses, further suggests that oxidative stress generated via ROS played a major role in initiating apoptosis [39]. This is consistent with the earlier DCFH-DA fluorescence results, where ZSS(0.15) produced the highest ROS levels, and with the photothermal data, where ZSS(0.15) showed the most efficient photothermal conversion. The enhanced efficacy of ZSS(0.15) over cisplatin highlights its potential as a non-invasive, laser-triggered alternative with lower systemic toxicity and higher selectivity toward cancer cells [40].

### 2.5. In-vivo Therapeutic Efficacy

#### 2.5.1. Animal model: Setup and treatment

To investigate the *in-vivo* phototherapeutic application, female athymic nude BALB/c mice were used to establish estrogen-supported MCF-7 human breast cancer xenografts (detailed experimental conditions are described in the *Supporting Information* file). Following tumor establishment (∼50 mm^3^; defined as Day 0; **Figure 5a**), mice were randomized into four groups (n=5 per group): (I) Control; (II) ZSS(0.15); (III) ZSS(0.15) +660 nm laser irradiation; and (IV) Cisplatin. ZSS(0.15) was administered via intratumoral injection at 10 mg kg^-1^ every 2 days. For the phototherapy group, tumors were irradiated 24 h post-injection using a 660 nm laser (0.5 W cm^-2^, 10 min), with irradiation conditions kept constant across animals (see *Experimental* Section for full protocol). Body weight (**Figure 5b**) in the control group increased steadily from 21.64Lg (Day 0) to 23.6Lg (Day 20), while ZSS(0.15)-treated mice showed a comparable rise from 21.7Lg to 22.7Lg. Notably, the ZSS(0.15)+660 nm group maintained stable weights (22.1Lg to 23.1Lg), indicating minimal systemic burden, unlike the cisplatin group, which showed a slight decline (21.5Lg to 21.9Lg), suggesting mild toxicity. Food intake (**Figure 5c**) further supported these observations: ZSS(0.15)+laser-treated mice consumed between 0.997-1.211Lg/day, slightly higher than control (0.797-1.168Lg/day), while cisplatin group showed a progressive decline after Day 10, dropping to 0.982-1.002Lg/day, indicative of treatment-associated distress. Similarly, water consumption (**Figure 5d**) in the ZSS(0.15)+laser group remained stable (16.1 to 14.3 mL), comparable to controls (16.4 to 14.5 mL), while cisplatin-treated mice exhibited a gradual decrease from 15.4 to 14.9 mL. In summary, these physiological metrics with stable body weight, sustained food and water intake, and absence of acute adverse effects strongly support the excellent biocompatibility of ZSS(0.15). More importantly, when irradiated with 660 nm irradiation, ZSS(0.15) maintained this biosafety profile, even while delivering potent photothermal and photodynamic tumor ablation, as established in prior *in-vitro* analyses. In contrast, cisplatin, although effective, exerted measurable systemic stress, evident from reduced food consumption and stagnant weight.

**Figure 5.**
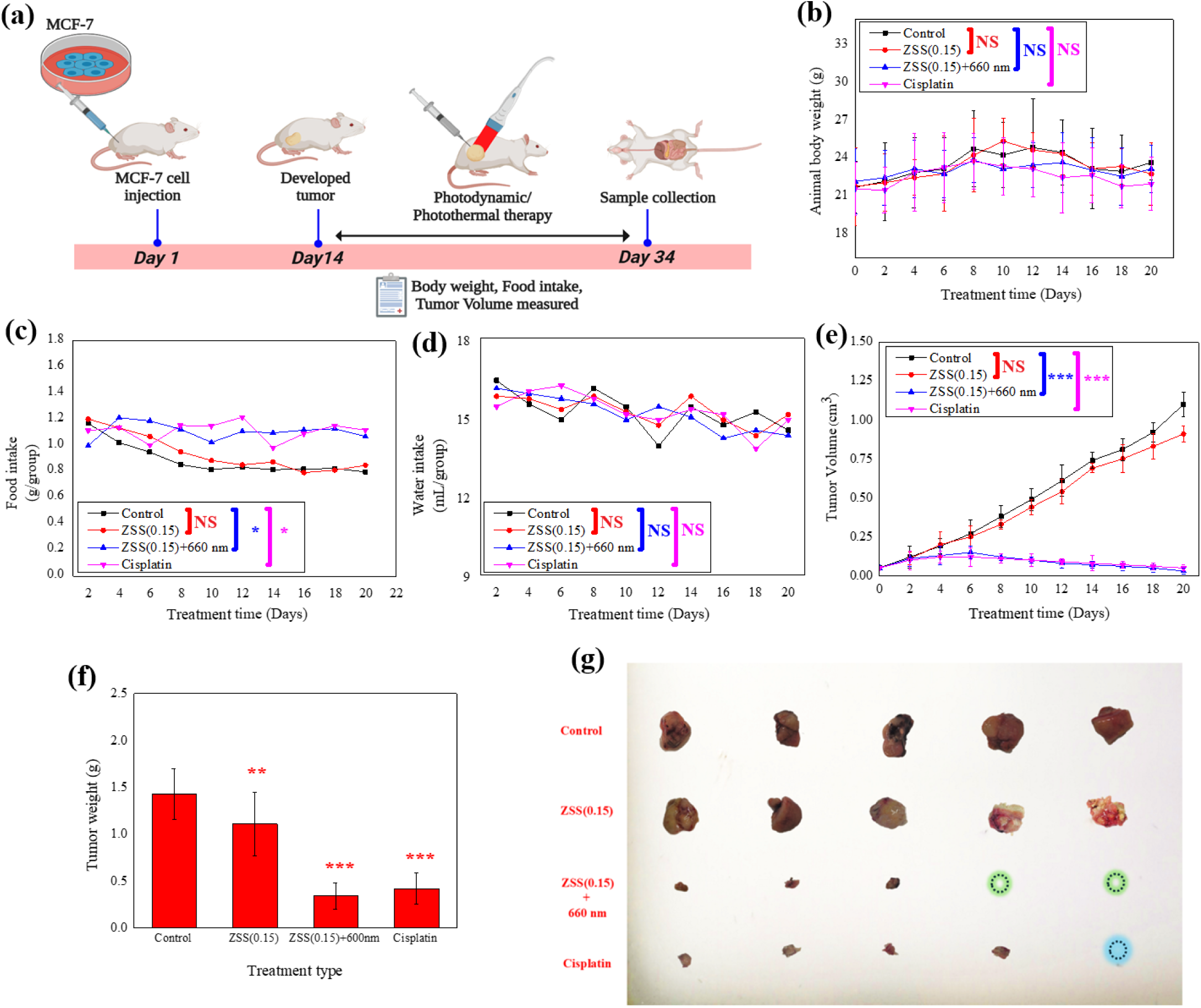
**(a)** Schematic timeline of animal treatment; **(b)** Body weight changes; **(c)** Daily food intake; **(d)** Daily water intake; **(e)** Tumor volume progression in different treatment groups; **(f)** Final tumor weight; **(g)** Representative images of excised tumors post-treatment. [Asterisks indicate statistically significant differences from control group; NS = no significance; * = *p* < 0.05; ** = *p* < 0.01; *** = *p* < 0.001].

#### 2.5.2. Tumor Suppression Efficacy and Visual Evidence

To evaluate the therapeutic efficacy of ZSS(0.15) *in-vivo*, tumor-bearing mice were treated over a 20-day period, and tumor progression was closely monitored. The control group exhibited rapid tumor growth (**Figure 5e**), with volumes escalating from 0.05 cm^3^ on Day 0 to 1.10 cm^3^ by Day 20, while the ZSS(0.15) - only group showed moderate suppression, reaching 0.91 cm^3^. Notably, the ZSS(0.15)+660 nm light-treated group demonstrated a dramatic reduction in tumor volume, declining to just 0.03 cm^3^ by Day 20, outperforming even the cisplatin group, which ended at 0.05 cm^3^. Tumor weight measurements (**Figure 5f**) corroborated these findings, ZSS(0.15)+light resulted in a significantly reduced tumor mass (0.34L±L0.14Lg), closely followed by cisplatin (0.42L±L0.17Lg), while ZSS(0.15) alone and control groups maintained higher tumor burdens of 1.11L±L0.34Lg and 1.43L±L0.27Lg, respectively. Representative tumor images (**Figure 5g**) revealed complete tumor ablation in 2 out of 5 mice in the ZSS(0.15)+660 nm group, and in 1 out of 5 mice in the cisplatin group, visually substantiating the quantitative findings. This superior tumor inhibition under light irradiation is directly linked to the photothermal and photodynamic potency of ZSS(0.15), as previously evidenced by its highest photothermal conversion efficiency and ROS generation capability, validated via photoluminescence quenching signals. Furthermore, enhanced charge separation and transfer properties observed through photocurrent and EIS studies suggest that ZSS(0.15) facilitates efficient electron/hole pair separation, crucial for ROS-mediated apoptosis [41]. The synergy of these properties culminates in profound tumor cell apoptosis and necrosis *in-vivo*. Collectively, these results firmly establish ZSS(0.15)+660 nm as a highly effective and biocompatible candidate for dual-modal photothermal and photodynamic cancer therapy [42], demonstrating tumor inhibition comparable to or even exceeding that of conventional chemotherapy, but with fewer systemic side effects.

#### 2.5.3. *Elisa*- Apoptotic marker (Pro/Anti) - Protein quantification

To further validate the apoptosis-inducing capabilities of ZSS(0.15), with and without 660 nm irradiation, the expression of key pro-apoptotic (Bax, Caspase-3, p53) and anti-apoptotic (Bcl-2, Survivin, PI3K, mTOR) proteins was quantified using ELISA in homogenized tumor tissues (Detailed experimental procedures are provided in the *Supporting Information* file) (**Figure 6**). The results revealed a significant upregulation of pro-apoptotic proteins (green bars) in the treatment groups, particularly in ZSS(0.15)+660 nm. Specifically, Bax expression rose from 42.3LJ±LJ3.4LJpg/mg in the control group to 68.9LJ±LJ5.7LJpg/mg in ZSS(0.15) and further to 102.6LJ±LJ8.4LJpg/mg in the ZSS(0.15)+660 nm group. Similarly, Caspase-3 levels increased from 36.5LJ±LJ2.7 to 82.1LJ±LJ6.4 and peaked at 145.5LJ±LJ11.7LJpg/mg. The tumor suppressor p53 [43] also followed this trend, increasing from 55.2LJ±LJ4.3 to 97.3LJ±LJ7.9 and then to 154.3LJ±LJ12.6LJpg/mg across the same groups. In contrast, anti-apoptotic proteins (red bars) were markedly suppressed in the treatment groups, most notably in the ZSS(0.15)+660 nm condition. Bcl-2 levels dropped from 111.9LJ±LJ9.2LJpg/mg in control to 63.6LJ±LJ5.3 in ZSS(0.15) and down to 32.8LJ±LJ2.9 in the irradiated group. Survivin, another potent anti-apoptotic marker [44], decreased from 89.4LJ±LJ7.2 to 47.5LJ±LJ4.2 and further to 24.2LJ±LJ1.8LJpg/mg, while PI3K and mTOR levels also exhibited a similar decline from 135.2LJ±LJ10.1 and 142.7LJ±LJ11.3 in the control group to 56.4LJ±LJ4.3 and 59.3LJ±LJ5.1LJpg/mg, respectively, in the ZSS(0.15)+660 nm group. Comparatively, the cisplatin group showed intermediate responses, with pro-apoptotic markers elevated (e.g., Caspase-3: 122.3LJ±LJ9.8LJpg/mg) and anti-apoptotic markers downregulated (e.g., Bcl-2: 40.9LJ±LJ3.9LJpg/mg), albeit less effectively than the laser-activated nanomaterial. These findings confirm that the phototherapeutic activation of ZSS(0.15) not only enhances apoptotic signaling but also robustly inhibits survival pathways, suggesting a dual mechanism involving mitochondrial-mediated apoptosis and PI3K/mTOR pathway suppression [45]. This is consistent with earlier photothermal and ROS studies showing high local hyperthermia and reactive oxygen generation under 660 nm irradiation. Together, these results strongly support the forthcoming mechanistic insights, emphasizing ZSS(0.15)+660 nm as a potent nano-therapeutic platform that disrupts tumor homeostasis at both the protein and signaling cascade levels.

**Figure 6.**
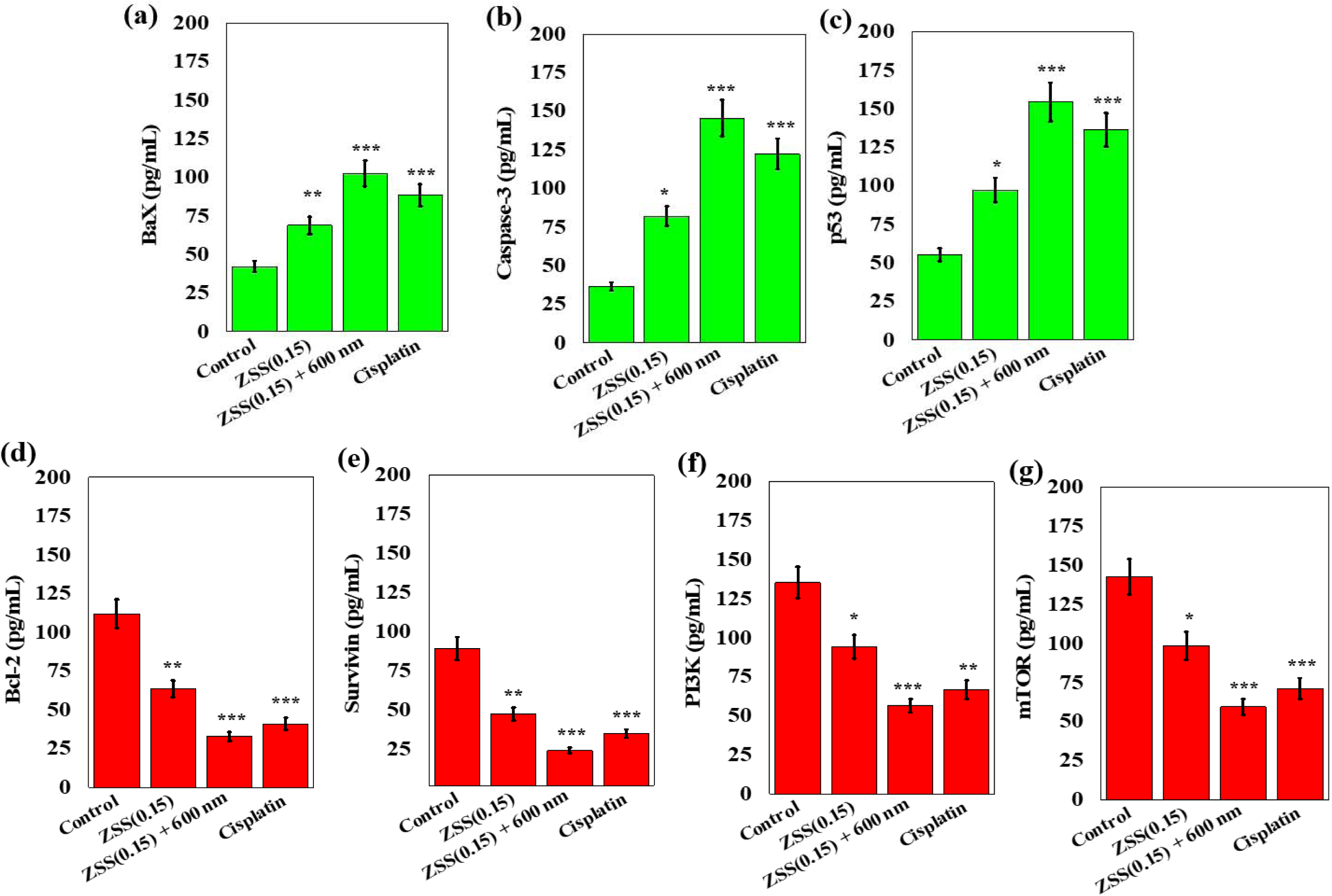
Protein quantification of apoptosis-related markers in tumor tissue homogenates using ELISA: **(a)** Bax, **(b)** Caspase-3, **(c)** p53, **(d)** Bcl-2, **(e)** Survivin, **(f)** PI3K, and **(g)** mTOR. [Asterisks indicate statistically significant differences from control group; NS = no significance; * = *p* < 0.05; ** = *p* < 0.01; *** = *p* < 0.001].

### 2.6. In-vivo Biosafety Evaluation

#### 2.6.1. Primary organ weight

To assess the systemic biosafety of ZSS(0.15)-based therapy, the primary organs: lung, liver, heart, spleen, and kidney were excised and weighed post-euthanasia (**Figure 7a**). Across all treatment groups, the organ weights remained largely comparable to the control group, indicating minimal systemic toxicity. Specifically, lung weights ranged from 0.67LJ±LJ0.15LJg in the ZSS(0.15)+660 nm group to 0.71LJ±LJ0.11LJg in the ZSS(0.15) group, closely matching the control (0.68LJ±LJ0.14LJg). Similarly, heart, spleen, and kidney weights showed only marginal fluctuations, with no statistically significant differences observed. For instance, kidney weights ranged from 0.80LJ±LJ0.20LJg to 0.89LJ±LJ0.20LJg, further supporting the absence of nephrotoxic effects. Notably, liver weight was slightly reduced in the ZSS(0.15)+660 nm group (2.01LJ±LJ0.30LJg) compared to control (2.40LJ±LJ0.20LJg), but this decrease was not accompanied by abnormal enlargement or discoloration typically indicative of hepatotoxicity. Visual examination (**Figure 7b**) revealed a subtle morphological difference in the livers of control mice (yellow arrow). However, the biological significance of this observation was not further evaluated, and no histological analyses were performed to determine its underlying cause. Importantly, no overt macroscopic abnormalities were observed in the treated groups. Collectively, the absence of significant changes in organ morphology, together with stable weight and physiological parameters, suggests that ZSS(0.15), particularly under 660 nm irradiation, did not induce observable organ-level toxicity under the present experimental conditions.

**Figure 7.**
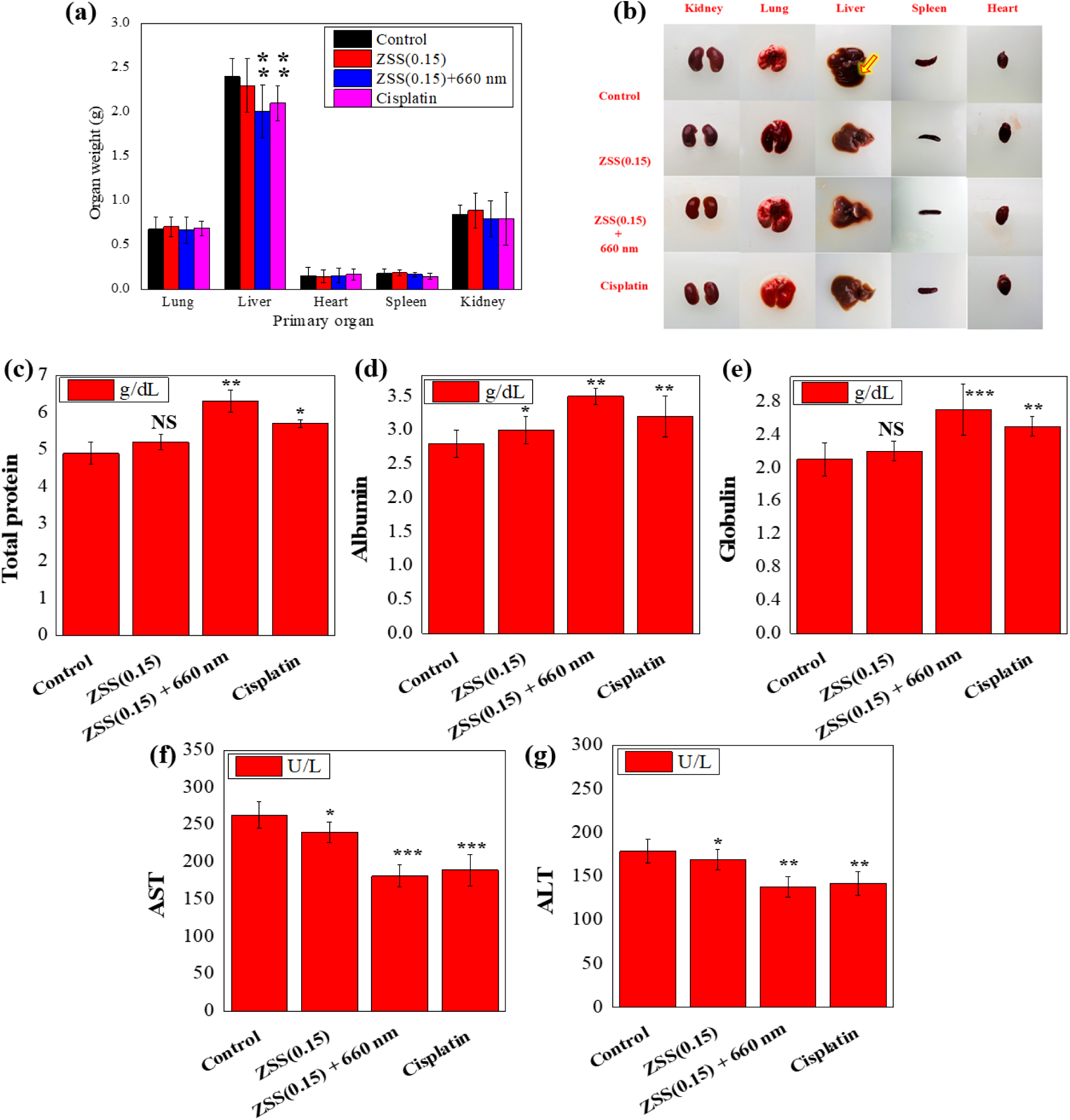
**(a)** Weights of primary organs and **(b)** their representative photographs; Liver function indicators measured by biochemical assays: **(c)** Total protein, **(d)** Albumin, **(e)** Globulin, **(f)** AST, and **(g)** ALT. [Asterisks indicate statistically significant differences from control group; NS = no significance; * = *p* < 0.05; ** = *p* < 0.01; *** = *p* < 0.001].

#### 2.6.2. Liver functionality analysis

To further validate the hepatic biosafety of the ZSS(0.15)-based therapeutic system, a comprehensive liver functionality assessment was conducted by quantifying serum levels of total protein, albumin, globulin, AST, and ALT (**Figure 7c-g**). Detailed tissue processing and assay procedures are provided in the *Supporting Information* file. The control group, burdened by unchecked tumor progression, exhibited mildly compromised liver function, reflected by lower protein values (total protein: 4.9LJ±LJ0.3LJg/dL; albumin: 2.8LJ±LJ0.2LJg/dL) and elevated transaminases (AST: 263LJ±LJ18LJU/L; ALT: 179LJ±LJ14LJU/L), suggesting hepatic stress or inflammation, consistent with the subtle liver bulging observed (**Figure 7b**). In stark contrast, the ZSS(0.15)+660 nm treated group demonstrated a markedly improved hepatic profile, with the highest levels of total protein (6.3LJ±LJ0.3LJg/dL), albumin (3.5LJ±LJ0.12LJg/dL), and globulin (2.7LJ±LJ0.31LJg/dL), coupled with significantly reduced AST (181LJ±LJ15LJU/L) and ALT (138LJ±LJ12LJU/L) levels, indicating robust liver function and minimal hepatic burden [46]. The standalone ZSS(0.15) and Cisplatin groups also showed moderate improvements over control but were inferior to the photodynamically enhanced ZSS(0.15)+660 nm treatment. These biochemical trends correlate well with the earlier primary organ weight data, where the liver in the ZSS(0.15)+660 nm group presented neither hypertrophy nor visible abnormalities, further reinforcing the biosafe nature of this nanoplatform. Thus, these findings not only dispel concerns of hepatic toxicity but also highlight the therapeutic efficiency of the ZSS(0.15)+660 nm system in mitigating systemic disease-induced organ stress, confirming its potential for safe clinical translation.

### 2.7. Mechanistic Insight

The therapeutic mechanism underlying the enhanced anticancer efficacy of ZSS(0.15) under 660 nm irradiation integrates both photothermal and photodynamic effects, synergistically promoting mitochondrial-mediated apoptosis in breast cancer cells. As depicted in the schematic (**Figure 8**), upon 660 nm light activation, ZSS(0.15) nanoflowers efficiently generate ROS alongside localized hyperthermia. These ROS act as a potent intracellular trigger, disrupting redox homeostasis and initiating mitochondrial dysfunction [47]. At the molecular level, ROS accumulation upregulates the tumor suppressor protein p53, which plays a pivotal role in promoting apoptosis. Elevated p53 expression facilitates the upregulation of the pro-apoptotic protein Bax while concurrently downregulating the anti-apoptotic protein Bcl-2, effectively tipping the balance toward cell death. This Bax/Bcl-2 modulation induces mitochondrial outer membrane permeabilization (MOMP), leading to the release of cytochrome c into the cytosol. Cytochrome c, once in the cytoplasm, binds to Apaf-1 and ATP to form the apoptosome, which subsequently activates initiator caspase-9 and executioner caspase-3 [48]. This cascade culminates in programmed cell death, as confirmed by significantly elevated caspase-3 levels in both *in-vitro* and *in-vivo* models treated with ZSS(0.15)+660 nm. Supporting the mechanistic cascade, *ELISA*-based protein quantification revealed markedly increased levels of Bax, Caspase-3, and p53, accompanied by strong suppression of anti-apoptotic markers such as Bcl-2 and Survivin. Moreover, key proliferative and survival-related signaling nodes: PI3K and mTOR, were significantly downregulated, indicating effective inhibition of the PI3K/Akt/mTOR survival axis [49]. These mechanistic insights are reinforced by the *in-vivo* tumor regression data, apoptotic gene expression profiles, and excellent biosafety evidenced by normal organ weights, preserved hepatic function, and non-toxic histological observations. Altogether, these findings validate that the ZSS(0.15)+660 nm system not only induces efficient apoptotic cell death through a ROS-mediated mitochondrial pathway but also suppresses pro-survival signaling making it a promising candidate for precise and safe combinatorial cancer therapy.

**Figure 8.**
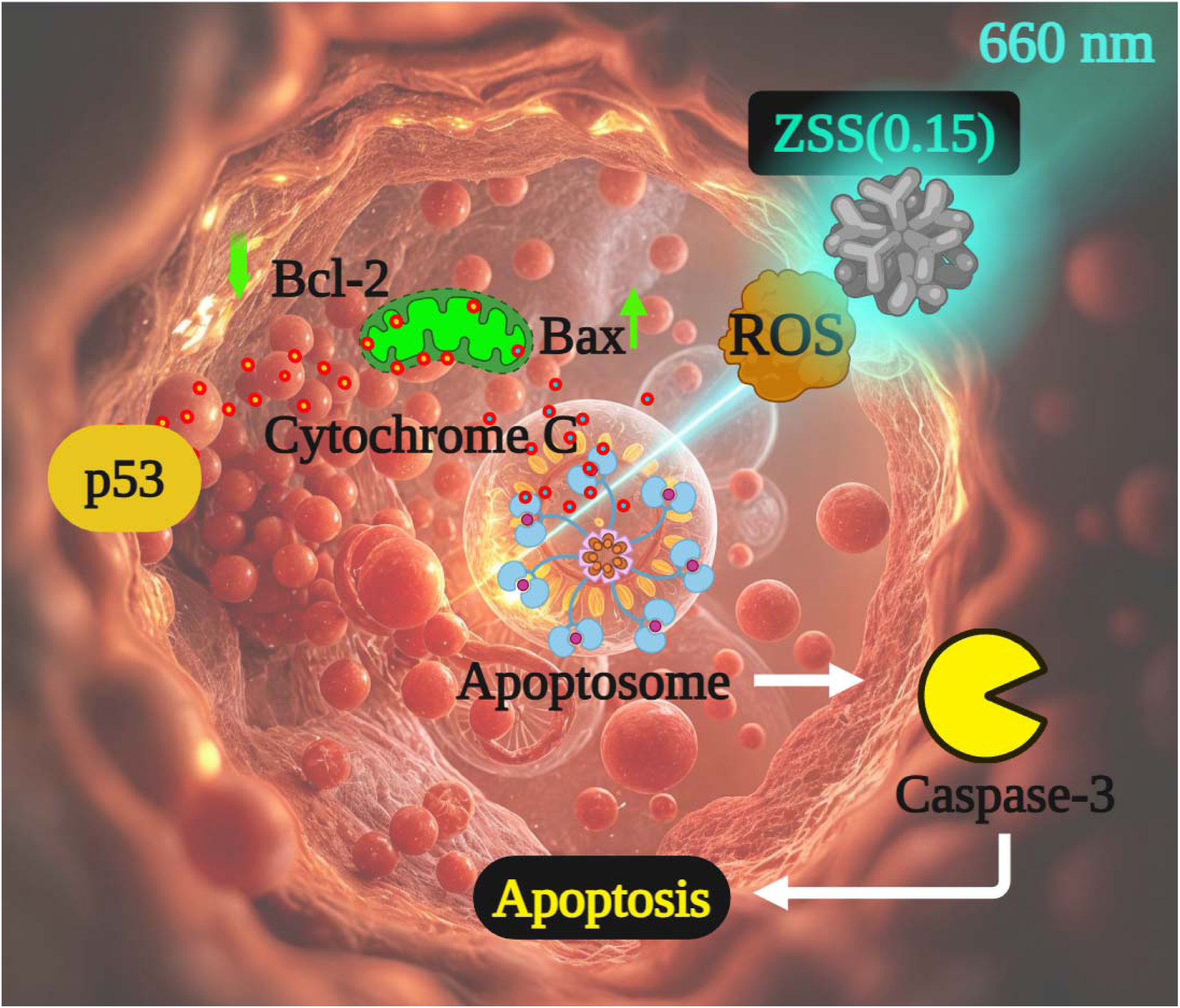
Proposed mechanism underlying the synergistic photothermal and photodynamic therapeutic effects of ZSS(0.15) against MCF-7 cells.

### 2.8. Broad-spectrum cytotoxicity of ZSS nanostructures across breast cancer subtypes

Breast cancer is a heterogeneous disease comprising molecularly distinct subtypes (luminal, HER2+, and triple-negative), each with different therapeutic vulnerabilities and resistance profiles [50]. Demonstrating efficacy across these subtypes is therefore critical for assessing the translational potential of new therapeutic platforms. Given the strong PTT/PDT effects of ZSS(0.15) against MCF-7 cells, we next evaluated its activity in triple-negative (MDA-MB-231), HER2+ (SK-BR-3), and luminal A (T47D) cells under identical conditions (660 nm, 0.5 W cm^-2^, 10 min). As shown in **Figure 9a**, all three lines exhibited a pronounced, concentration-dependent decrease in viability, with IC_50_ values of 16.8 ± 0.9 μg/mL (MDA-MB-231), 15.6 ± 0.7 μg/mL (SK-BR-3), and 15.9 ± 0.8 μg/mL (T47D), indicating largely preserved potency across molecular subtypes. At the IC_50_ concentration for MCF-7 cells (15 μg/mL), SK-BR-3 and T47D showed viability close to 50%, whereas MDA-MB-231 retained ∼60% viability (**Figure 9b**), consistent with the more aggressive and treatment-resistant phenotype typically associated with triple-negative breast cancers [51]. Importantly, these results demonstrate that Zn-doped AgLJS nanostructures maintain consistent phototherapeutic potency across ER+, HER2+, and triple-negative breast cancer cell lines, with comparable ICLJLJ values observed among the different molecular subtypes. The broad-spectrum efficacy further supports the notion that Zn-induced enhancements in photothermal conversion and ROS generation are subtype-independent, providing a versatile nanoplatform for diverse clinical settings. While *in-vivo* validation in additional tumor models will be required to fully establish subtype-independent therapeutic efficacy, the consistent phototherapeutic response observed across multiple molecular subtypes *in-vitro* strongly suggests that the underlying PTT/PDT mechanism of ZSS(0.15) is not restricted to a single breast cancer phenotype.

**Figure 9.**
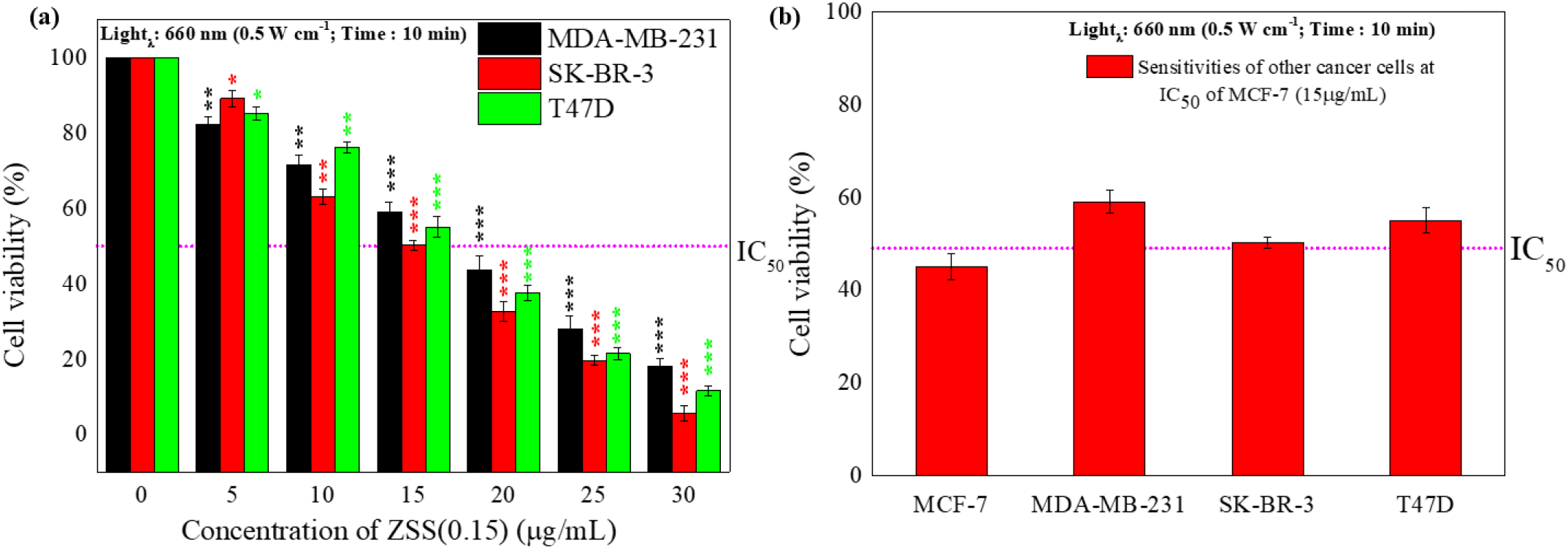
**(a)** Cell viability (MTT assay) under combined photothermal and photodynamic treatment (PTT/PDT) of ZSS(0.15) against other selected cancer cell lines [Asterisks indicate statistically significant differences from 0 µg/mL within each treatment group; NS = no significance; * = *p* < 0.05; ** = *p* < 0.01; *** = *p* < 0.001] **(b)** comparative plot of IC_50_ values for the select cancer cell lines.

## 3. CONCLUSION

In summary, we developed Zn-doped Ag_2_S nanostructures (ZSS), with ZSS(0.15) identified as the most effective composition for light-triggered cancer therapy. Under 660 nm irradiation, this nanoplatform combined strong photothermal heating with efficient ROS generation, enabling synergistic therapeutic effects.

*Key outcomes of this work include*:

- **Enhanced therapeutic efficacy**: ZSS(0.15) achieved nearly 97% tumor suppression *in-vivo* with no systemic toxicity.
- **Mechanistic insight**: Zn doping favored improved charge separation and light utilization, boosting both photothermal and photodynamic activity.
- **Safety advantage**: Compared to cisplatin, ZSS(0.15) showed minimal side effects while maintaining superior cancer cell killing.
- **Broad applicability**: Consistent performance was demonstrated across multiple breast cancer subtypes, not limited to a single model.

These findings highlight Zn doping as a powerful strategy to optimize Ag_2_S-based nanoplatforms for dual-mode cancer therapy. While highly encouraging, further evaluation in orthotopic and metastatic tumor models, integration with targeted delivery systems, and exploration of deeper tissue-penetrating light sources will be critical for clinical translation. Taken together, our study positions ZSS(0.15) as a promising next-generation nanomedicine for safe, effective, and versatile photothermal-photodynamic breast cancer therapy.

## 4. EXPERIMENTAL

### 4.1. Materials

All reagents and chemicals employed in this study were purchased from Sigma-Aldrich (South Korea) and used without any further purification. Deionized water (DI water) was used in all procedures involving aqueous media.

### 4.2. Synthesis of Zn_x_Ag_2-2x_S (ZSS)nanostructures

The synthesis of Zn_x_Ag_2-2x_S nanostructures (ZSS) was accomplished via a two-step hydrothermal method involving a cation exchange strategy. Initially, pristine Ag_2_S nanostructures were synthesized by dissolving silver nitrate (AgNO_3_, 0.02 M) in 50 mL of deionized water and stirring at 25 °C for 30 minutes at 600 rpm. A freshly prepared aqueous solution of sodium sulfide (Na_2_S, 0.01 M, 50 mL) was then added dropwise under constant stirring (600 rpm), resulting in a black dispersion, which was stirred further for 6 hours at room temperature. The mixture was then transferred to a Teflon-lined autoclave and subjected to hydrothermal treatment at 150 °C for 16 hours. After natural cooling to room temperature, the Ag_2_S product was collected, washed thoroughly with deionized water and ethanol, and subsequently dried. To obtain the ZSS nanostructures, the as-synthesized Ag_2_S was redispersed in deionized water and treated with varying concentrations of zinc nitrate (Zn(NO_3_)_2_) solution to achieve controlled Ag^+^/Zn^2+^ ion exchange. The molar ratios of Ag^+^:Zn^2+^ were adjusted as 1.95:0.05, 1.90:0.10, 1.85:0.15, 1.80:0.20, 1.75:0.25, and 1.70:0.30. The mixture was stirred at 25 °C for 30 minutes and then subjected to a second hydrothermal treatment at 150 °C for 16 hours to facilitate partial substitution of Ag^+^ by Zn^2+^ within the Ag_2_S lattice. The resulting Zn_x_Ag_2-2x_S nanostructures were washed with deionized water and ethanol, followed by freeze-drying. For comparison, pure ZnS nanostructures were synthesized under identical hydrothermal conditions by replacing AgNO_3_ with Zn(NO_3_)_2_ (**Figure 1a**).

### 4.3. In-vitro studies

#### 4.3.1. MTT assay

Breast cancer cell line culture conditions, maintenance protocols, and seeding procedures were performed following standard methods, as detailed in the *Supporting Information* file. MCF-7 cells were seeded into 96-well plates at a density of 5 × 10^3^ cells per well and incubated for 24 hours to allow for attachment. After incubation, the culture medium was replaced with 100 µL of fresh DMEM containing various concentrations of the synthesized nanostructures (5, 10, 15, 20, and 25 µg/mL). For PDT evaluation, cells were exposed to a 660 nm laser at an intensity of 0.5 W/cm^2^ for 10 minutes, followed by incubation for an additional 24 hours. The laser source was mounted on a fixed stand at a constant distance (∼6 cm) above the culture plate to ensure uniform and reproducible irradiation conditions. During irradiation, culture plates were placed inside a sterile biosafety cabinet and exposed to 660 nm light at room temperature for 10 min with the plate lid temporarily removed to ensure uniform light delivery. Cells remained in complete culture medium during exposure. Immediately after irradiation, plates were returned to a 37 °C, 5% CO_2_ incubator for further incubation. The cells in control groups were kept in the dark under identical conditions. Subsequently, 10 µL of MTT solution (5 mg/mL in PBS) was added to each well, and the plates were incubated for 4 hours at 37 °C to allow the formation of formazan crystals. The medium was then carefully removed, and 100 µL of dimethyl sulfoxide (DMSO) was added to solubilize the crystals. Absorbance was measured at 570 nm using a microplate reader (Infinite M1000 Pro, Tecan, Switzerland). Untreated cells were used as the negative control, while cells treated with cisplatin served as the positive control [52].

#### 4.3.2. FACS

To evaluate apoptosis and necrosis induced by ZSS nanostructures, cells were stained with Annexin V-FITC and propidium iodide (PI) following the manufacturer’s instructions. In brief, cells were seeded into 6-well plates at a density of 2 × 10^5^ cells per well and incubated for 24 hours to facilitate attachment. After incubation, the culture medium was replaced with fresh medium containing ZSS nanostructures. Cells were then irradiated with 660 nm red light at a power density of 0.5 W/cm^2^ for 10 minutes under sterile conditions inside a biosafety cabinet. The laser source was mounted on a fixed stand at a constant distance (∼6 cm) above the culture plate to ensure uniform and reproducible irradiation conditions. During irradiation, cells remained in complete medium at room temperature with the plate lid temporarily removed to ensure uniform light exposure. Immediately after irradiation, plates were returned to a 37 °C, 5% CO_2_ incubator for further incubation. Following that 24-hour incubation, cells were harvested, rinsed twice with cold phosphate-buffered saline (PBS), and stained with Annexin V-FITC and PI. Staining was performed in the dark for 15 minutes at room temperature to minimize light-induced degradation. The stained cells were then analyzed by flow cytometry using a BD Accuri C6 cytometer to quantify the proportions of viable, early apoptotic, late apoptotic, and necrotic cells [53].

### 4.4. In-vivo studies

All animal experiments were conducted in accordance with institutional ethical guidelines and were approved by the Institutional Animal Care and Use Committee (IACUC). Detailed animal housing, care conditions, and ethical compliance information are provided in the *Supporting Information* file.

#### 4.4.1. Animal treatment

The MCF-7 xenograft tumor model was established using standard estrogen-supported implantation procedures. Detailed tumor induction protocols and growth monitoring methods are described in the *Supporting Information* file. When the tumors reached an approximate volume of 50 mm^3^, the mice were randomly divided into four experimental groups (nLJ=LJ5 per group). Therapeutic interventions were carried out over a 20-day treatment period. ZSS(0.15) was prepared in sterile PBS at a concentration of 4 mg/mL and administered via intratumoral injection at a dose of 10 mg/kg body weight. The injection volume was adjusted according to individual animal body weight and ranged between 40 to 80 µL per mouse to ensure consistent local delivery. Injections were performed once every two days. Cisplatin was administered under the same schedule at an equivalent dose. For the groups receiving photothermal therapy, 660 nm red-light laser irradiation (power density: 0.5 W/cm^2^, Irradiation time: 10 min) was applied 24 hours after each intratumoral injection to ensure uniform local distribution of the nanostructures within the tumor prior to light exposure. Laser irradiation (660 nm, 0.5 W cm^-2^, 10 min) was performed using a fixed irradiation platform. During irradiation, mice were placed in a ventilated restraining enclosure designed to minimize movement while avoiding excessive stress. Anesthesia was not applied, as the procedures were brief and minimally invasive. Tumors were positioned perpendicular to the laser beam to ensure uniform illumination. The irradiation distance and spot size were kept constant for all animals, and laser output was verified prior to each session to maintain a consistent power density across treated tumors. Tumor dimensions were measured every two days using a vernier caliper, and tumor volume was calculated. Additionally, body weight, food intake, and water consumption were recorded to evaluate the general health and treatment response of the animals. At the end of the study, all mice were humanely euthanized using carbon dioxide asphyxiation. Tumors were excised, weighed, and preserved for further analyses. Major organs, including the kidneys, lungs, liver, spleen, and heart were also collected and weighed for subsequent assessments.

### 4.5. Broad-Spectrum Cytotoxicity Across Breast Cancer Subtypes

While MCF-7 cells were selected as the primary model to evaluate the mechanistic basis of ZSS-mediated photothermal and photodynamic effects, we further extended the investigation to additional breast cancer subtypes (MDA-MB-231; SK-BR-3; T47D) cells under identical conditions (ZSS(0.15, 660 nm, 0.5 W cm^-2^, 10 min) to assess the broader applicability of the treatment.

### 4.6. Statistical analysis

All statistical analyses were conducted using SPSS software (version 22). Differences among multiple groups were assessed by one-way analysis of variance (ANOVA), followed by Tukey’s post hoc test to determine specific group differences. *p*-value < 0.05 was considered statistically significant. All the results are presented as Average ± standard deviation (SD).

## SUPPORTING INFORMATION

Supporting Information is available from the Wiley Online Library or from the author.

## ACKNOWLEDGEMENTS

This work was supported by the National Research Foundation of Korea (NRF) grant funded by the Korea government, MSIT [RS-2025-00523589] and MOE [2020R1I1A3054429], and by research funds from Jeonbuk National University in 2023. The authors acknowledge the use of the *NightCafe* AI image generator for background artwork in the *Graphical abstract* and *Mechanistic Insight* figure; all scientific annotations were created and verified by the authors.

## CONFLICTS OF INTEREST

The authors declare that they have no known competing financial interests or personal relationships that could have influenced the findings reported in this paper.

## DATA AVAILABILITY

Relevant data can be made available on request.

Received: ((will be filled in by the editorial staff))

Revised: ((will be filled in by the editorial staff))

Published online: ((will be filled in by the editorial staff))

## Notes

### Competing Interest Statement

The authors have declared no competing interest.

### Summary of Updates

In response to the reviewers suggestions, we have performed additional experiments and clarifications, including: 1.Photothermal stability evaluation using repeated laser ON/OFF cycling. 2.Identification and quantitative assessment of singlet oxygen generation using SOSG and DPBF photobleaching kinetics. 3.Clarification and correction of in-vivo experimental parameters, including intratumoral administration route, dosage, and irradiation conditions. 4.Serum stability analysis under physiologically relevant conditions, provided in the Supporting Information file. 5.Standardization of the in-vivo xenograft model description (mouse strain and tumor establishment details), now stated consistently across the manuscript and Supporting Information. In addition, terminology inconsistencies and interpretational ambiguities identified by the reviewers, such as irradiation wavelength classification, bandgap description, and the distinction between structural optimization and therapeutic performance have been carefully corrected throughout the manuscript.

## REFERENCES

1. Kim, J.; Harper, A.; McCormack, V.; Sung, H.; Houssami, N.; Morgan, E.; Mutebi, M.; Garvey, G.; Soerjomataram, I.; Fidler-Benaoudia, M. M., Nature Medicine 2025, 1–9.

2. Sonkin, D.; Thomas, A.; Teicher, B. A., Cancer Genetics 2024, 286-287, 18–24. DOI 10.1016/j.cancergen.2024.06.002.

3. Xiong, X.; Zheng, L.-W.; Ding, Y.; Chen, Y.-F.; Cai, Y.-W.; Wang, L.-P.; Huang, L.; Liu, C.-C.; Shao, Z.-M.; Yu, K.-D., Signal Transduction and Targeted Therapy 2025, 10 (1), 49.

4. Nkune, N. W.; Abrahamse, H., Journal of Biophotonics 2025, n/a (n/a), e70005. DOI 10.1002/jbio.70005.

5. Zhao, Z.; Chen, T.; Li, J.; Xue, X.; Ge, J.; Wang, P., Smart Molecules 2025, 3 (2), e20240049. DOI 10.1002/smo.20240049.

6. Yang, F.; Chen, F.; Li, S.; Wang, S.; Peng, L.; Deng, S.; Zhang, F.; Jeschke, U.; Wang, Y.; Luo, C., Chemical Engineering Journal 2025, 516, 163962. DOI 10.1016/j.cej.2025.163962.

7. Papa, V.; Furci, F.; Minciullo, P. L.; Casciaro, M.; Allegra, A.; Gangemi, S. Photodynamic Therapy in Cancer: Insights into Cellular and Molecular Pathways Current Issues in Molecular Biology [Online], 2025.

8. Jyothish, B.; Jacob, J., Surfaces and Interfaces 2025, 56, 105553. DOI 10.1016/j.surfin.2024.105553.

9. Hu, J.-J.; Chen, Y.; Lou, X.; Xia, F.; Wu, X.; Yoon, J., Coordination Chemistry Reviews 2025, 532, 216526. DOI 10.1016/j.ccr.2025.216526.

10. Lin, Y.; Han, D.; Li, Y.; Tan, L.; Liu, X.; Cui, Z.; Yang, X.; Li, Z.; Liang, Y.; Zhu, S.; Wu, S., ACS Sustainable Chemistry & Engineering 2019, 7 (17), 14982–14990. DOI 10.1021/acssuschemeng.9b03287.

11. Zhu, K.; Li, Z.; Cao, J.; Cao, Y.; Wang, J.; Wang, S.; Chen, L.; Zhou, H.; Huang, W.; Zou, H.; Li, Q.; Mu, J.; Song, J., Advanced Science 2025, n/a (n/a), 2417828. DOI 10.1002/advs.202417828.

12. Oh, H. S.; Mohan, H.; Sathya, P. M.; Kim, G.; Ha, G. H.; Shin, T., Materials Today Communications 2023, 34, 105194.

13. Zhang, Y.; Zhang, Y.; Li, Y.; Fu, Y.; Zhao, Y.; Zhao, W.; Li, R.; Xian, Y.; Tu, K.; Wu, F.; Li, C.; Hou, Y.; Zhang, M., Chemical Engineering Journal 2023, 474, 145685. DOI 10.1016/j.cej.2023.145685.

14. Wang, J.; Huang, Z.; Wu, Y.; Jiang, X.; Ji, Y.; Braeckmans, K.; Wang, M.; Wang, L.; Chen, W. R.; Xia, Y.; Tang, Z.; Xu, X., BMC Biology 2025, 23 (1), 111. DOI 10.1186/s12915-025-02215-w.

15. Liu, T.; Zhang, X.; Liu, D.; Chen, B.; Ge, X.; Gao, S.; Song, J., Advanced Optical Materials 2021, 9 (12), 2100233. DOI 10.1002/adom.202100233.

16. Tabrizi, N.; Jamali-Sheini, F.; Ebrahimiasl, S.; Cheraghizade, M., Optical Materials 2024, 152, 115374. DOI 10.1016/j.optmat.2024.115374.

17. Wang, C.; Zheng, H.; Ma, R.; Zheng, X.; Guan, X. Ag2S/Zn2+ -Decorated g-C3N4 Type-II Heterojunction with Wide-Spectrum Response: Construction and Photocatalytic Performance in Ciprofloxacin Degradation Molecules [Online], 2025.

18 Tang, Z.; Yang, H.; Sun, Z.; Zhang, Y.; Chen, G.; Wang, Q., Nano Research 2023, 16 (10), 12315–12322. DOI 10.1007/s12274-023-5952-z.

19. Dai, X.; Cheng, H.; Bai, Z.; Li, J., Journal of Cancer 2017, 8 (16), 3131.

20. Orrantia-Borunda, E.; Anchondo-Nuñez, P.; Acuña-Aguilar, L. E.; Gómez-Valles, F. O.; Ramírez-Valdespino, C. A., Breast Cancer [Internet] 2022.

21. Mandari, K. K.; Im, Y.; Lee, Y.-A.; Pandey, S.; Kang, M., Applied Surface Science 2025, 704, 163452. DOI 10.1016/j.apsusc.2025.163452.

22. Patil, B. N., Journal of Applied Spectroscopy 2025, 92 (1), 146–150. DOI 10.1007/s10812-025-01890-5.

23. Yin Win, K.; Feng, S.-S., Biomaterials 2005, 26 (15), 2713–2722. DOI 10.1016/j.biomaterials.2004.07.050.

24. Feng, H.; Wang, J.; Fan, W.; Zhang, C., Materials Letters 2014, 126, 67–70. DOI 10.1016/j.matlet.2014.04.037.

25. Geravand, M.; Jamali-Sheini, F., Advanced Powder Technology 2019, 30 (2), 347–358. DOI 10.1016/j.apt.2018.11.012.

26. Yang, Z.-C.; Gu, Q.-S.; Chao, J.-J.; Tan, F.-Y.; Mao, G.-J.; Hu, L.; Ouyang, J.; Li, C.-Y., Analytica Chimica Acta 2024, 1316, 342860. DOI 10.1016/j.aca.2024.342860.

27. Jagadeeswararao, M.; Swarnkar, A.; Markad, G. B.; Nag, A., The Journal of Physical Chemistry C 2016, 120 (34), 19461–19469. DOI 10.1021/acs.jpcc.6b06394.

28. Zhang, Z.; Yan, H.; Qiu, B.; Ran, P.; Cao, W.; Jia, X.; Huang, K.; Li, X., Small 2022, 18 (21), 2200813. DOI 10.1002/smll.202200813.

29. Guo, M.; Xing, Z.; Zhao, T.; Li, Z.; Yang, S.; Zhou, W., Applied Catalysis B: Environmental 2019, 257, 117913. DOI 10.1016/j.apcatb.2019.117913.

30. Chen, Q.; Yang, X.; Fu, X.; Pan, J.; Zhou, J.; Zhou, G.; Cheng, K., Chemical Engineering Journal 2024, 499, 156242. DOI 10.1016/j.cej.2024.156242.

31. Mohan, H.; Ramalingam, V.; Adithan, A.; Natesan, K.; Seralathan, K.-K.; Shin, T., Journal of Hazardous Materials 2021, 416, 126209.

32. Tiwari, N.; Gaikwad, P.; Kamat, R. K.; Kulkarni, S., Nano Trends 2025, 10, 100110. DOI 10.1016/j.nwnano.2025.100110.

33. Shou, P.; Yu, Z.; Wu, Y.; Feng, Q.; Zhou, B.; Xing, J.; Liu, C.; Tu, J.; Akakuru, O. U.; Ye, Z.; Zhang, X.; Lu, Z.; Zhang, L.; Wu, A., Advanced Healthcare Materials 2020, 9 (1), 1900948. DOI 10.1002/adhm.201900948.

34. Vinícius-Araújo, M.; Shrivastava, N.; Sousa-Junior, A. A.; Mendanha, S. A.; Santana, R. C. D.; Bakuzis, A. F., ACS Applied Nano Materials 2021, 4 (2), 2190–2210. DOI 10.1021/acsanm.1c00027.

35. Almuqbil, R. M., Advances in Pharmacological and Pharmaceutical Sciences 2024, 2024 (1), 4646855. DOI 10.1155/2024/4646855.

36. Solanki, R.; Rajput, P. K.; Jodha, B.; Yadav, U. C. S.; Patel, S., Scientific Reports 2024, 14 (1), 2595. DOI 10.1038/s41598-024-51970-3.

37. Kong, L.; Xing, L.; Zhou, B.; Du, L.; Shi, X., ACS Applied Materials & Interfaces 2017, 9 (19), 15995–16005. DOI 10.1021/acsami.7b03371.

38. Li, D.; Zhang, Y.; Wen, S.; Song, Y.; Tang, Y.; Zhu, X.; Shen, M.; Mignani, S.; Majoral, J.-P.; Zhao, Q., Journal of Materials Chemistry B 2016, 4 (23), 4216–4226.

39. Li, S.; Lui, K.-H.; Lau, W.-S.; Chen, J.; Lo, W.-S.; Li, X.; Gu, Y.-J.; Lin, L.-t.; Wong, W.-T., ACS Applied Materials & Interfaces 2022, 14 (29), 33712–33725. DOI 10.1021/acsami.2c07592.

40. Qi, M.-H.; Wang, D.-D.; Qian, W.; Zhang, Z.-L.; Ao, Y.-W.; Li, J.-M.; Huang, S.- W., Advanced Materials 2025, 37 (5), 2412191. DOI 10.1002/adma.202412191.

41. Qi, B.; Xiao, Z.; Huang, J.; Zhou, G.; Zhou, T.; Liu, G.; Li, Z., Chemical Engineering Journal 2024, 498, 155432. DOI 10.1016/j.cej.2024.155432.

42. Xu, M.-Y.; Zeng, J.; Han, X.-H.; Wei, D.-D.; Fan, B.; Qu, X. H.; Huang, S.; Han, X.- J., Materials & Design 2025, 251, 113647. DOI 10.1016/j.matdes.2025.113647.

43. Jiang, Y.; Li, Y.; Wang, Y.; Li, X., Biochemical Pharmacology 2024, 227, 116456. DOI 10.1016/j.bcp.2024.116456.

44. Zhang, L.; Yi, B.; Chen, J., Arabian Journal of Chemistry 2024, 17 (9), 105861. DOI 10.1016/j.arabjc.2024.105861.

45. Wang, Z.; Guo, Y.; Li, K.; Huo, Y.; Wang, S.; Dong, S.; Ma, M., Bioorganic & Medicinal Chemistry 2024, 115, 117908. DOI 10.1016/j.bmc.2024.117908.

46. Yu, B.; Liu, M.; Jiang, L.; Xu, C.; Hu, H.; Huang, T.; Xu, D.; Wang, N.; Li, Q.; Tang, B. Z.; Huang, X.; Zhang, W., Advanced Healthcare Materials 2024, 13 (11), 2303643. DOI 10.1002/adhm.202303643.

47. Jin, L.; Zhou, S.; Zhang, T.; Cui, F.; Yu, H.; Wang, J.; Wu, M.; Wang, Y.; Song, S.; Zhang, S., Small 2025, 21 (2), 2408639.

48. Rajabzadeh, P.; Zoorazma, P.; Abbasi, M.; Sadat Shandiz, S. A., Scientific Reports 2025, 15 (1), 1–16.

49. Chang, Y.; Shen, S.; Zhang, L.; Zeng, J.; Sun, J.; Guo, Y.-W.; Su, M.-z., Chemistry & Biodiversity, e202500114.

50. Carvalho, E.; Canberk, S.; Schmitt, F.; Vale, N., Cancers 2025, 17 (7), 1102.

51. Arora, R.; Yates, C.; Gary, B. D.; McClellan, S.; Tan, M.; Xi, Y.; Reed, E.; Piazza, G. A.; Owen, L. B.; Dean-Colomb, W., PLoS One 2014, 9 (6), e98370.

52. Li, R.-T.; Zhu, Y.-D.; Li, W.-Y.; Hou, Y.-K.; Zou, Y.-M.; Zhao, Y.-H.; Zou, Q.; Zhang, W.-H.; Chen, J.-X., Journal of Nanobiotechnology 2022, 20 (1), 212. DOI 10.1186/s12951-022-01427-4.

53. Mohan, H.; Ramalingam, V.; Lim, J.-M.; Lee, S.-W.; Kim, J.; Lee, J.-H.; Park, Y.-J.; Seralathan, K.-K.; Oh, B.-T., Colloids and Surfaces A: Physicochemical and Engineering Aspects 2020, 607, 125469. DOI 10.1016/j.colsurfa.2020.125469.

