## Supplementary material for "Cation-exchange synthesized Zn-doped Ag_2_S Nanostructures for Photothermal and Photodynamic Therapies across Breast Cancer Subtypes": Figure SF1 & 2; Table ST1 to 3

***Additional Information file for,***

Satabdi Acharya

Department of Medical Science, Jeonbuk National University, Jeonju 54896, Republic of Korea.

Inhee Chung **(****)**

Department of Anatomy and Cell Biology, School of Medicine and Health Sciences, George Washington University, Washington, DC, USA

**Short title:** *Zn_x_Ag_2-2x_S (ZSS) for efficient photodynamic/thermal activity*

**This file contains,**

**METHODS**

**1.	*In-vitro* studies**

**1.1.	Cell culture**

**1.2.	Fluorescent staining**

**1.3.	Gene expression analysis**

**2.	*In-vivo* studies**

**2.1.	Animal care and Ethical Compliance**

**2.2.	Tumor induction**

**2.3. *Elisa*- Apoptotic marker (Pro/Anti) - Protein quantification**

**2.4.	Liver functionality test**

**TABLES**

**Table ST1** List of chemicals, kits, and cell lines used in the study

**Table ST2** Primer information for RT-PCR analysis

**Table ST3** Charge transfer resistance (Rct) values obtained from equivalent circuit fitting of EIS data.

**FIGURES**

**Figure SF1** Particle size distribution of ZSS(0.15)

**Figure SF2** Real-time temperature profile and photothermal conversion efficiency

**Figure SF3** **(a)** Time-dependent changes in hydrodynamic size measured by dynamic light scattering (DLS) for ZSS(0.15) dispersed in PBS (pH 7.4) and PBS containing 10% fetal bovine serum (FBS); **(b)** Corresponding polydispersity index (PDI) and **(c)** ζ-potential of ZSS(0.15) measured at 0 h and 24 h in PBS and PBS + 10% FBS

**Table ST1:** List of chemicals, kits, and cell lines used in the study

| **S. No** | **Chemical name** | **CAS No** | **Source** | **City** | **Country** |
| --- | --- | --- | --- | --- | --- |
| 1 | Silver nitrate | 209139 | Sigma Aldrich | Seoul | South Korea |
| 2 | Sodium sulfide | 431648 | Sigma Aldrich | Seoul | South Korea |
| 3 | Zinc nitrate | 96482 | Sigma Aldrich | Seoul | South Korea |
| 4 | 3-(4,5-Dimethyl-2-thiazolyl)-2,5-diphenyl-2H-tetrazolium Bromide (MTT) | 475989 | Sigma Aldrich | Seoul | South Korea |
| 5 | Phosphate buffered saline (PBS) | P2272 | Sigma Aldrich | Seoul | South Korea |
| 6 | Cisplatin | 232120 | Sigma Aldrich | Seoul | South Korea |
| 7 | Dimethyl sulfoxide (DMSO) | 472301 | Sigma Aldrich | Seoul | South Korea |
| 8 | Dulbecco′s Modified Eagle′s Medium (DMEM medium) | D6429 | Sigma Aldrich | Seoul | South Korea |
| 9 | Fetal Bovine Serum (FBS) | F0850 | Sigma Aldrich | Seoul | South Korea |
| 10 | Penicillin-streptomycin solution | P7539 | Sigma Aldrich | Seoul | South Korea |
| 11 | Paraformaldehyde | P6148 | Sigma Aldrich | Seoul | South Korea |
| 12 | Hydrogen Peroxide | H1009 | Sigma Aldrich | Seoul | South Korea |
| 13 | Dihydroethidium | 37291 | Sigma Aldrich | Seoul | South Korea |
| 14 | DAPI | D9542 | Sigma Aldrich | Seoul | South Korea |
| 15 | Annexin-V FITC and propidium iodide | APOAF | Sigma Aldrich | Seoul | South Korea |
| 16 | TRIzol reagent | 15596026 | Thermo Fisher Scientific | Seoul | South Korea |
| 17 | BCL2 | - | TaKaRa | Seoul | South Korea |
| 18 | BAX | - | TaKaRa | Seoul | South Korea |
| 19 | P53 | - | TaKaRa | Seoul | South Korea |
| 20 | CASP3 | - | TaKaRa | Seoul | South Korea |
| 21 | GADPH | - | TaKaRa | Seoul | South Korea |
| 22 | *Bax* | MBS2513810 | MyBiosource | San Diego, CA | USA |
| 23 | *Caspase 3* | MBS260710 | MyBiosource | San Diego, CA | USA |
| 24 | *P53* | MBS355295 | MyBiosource | San Diego, CA | USA |
| 25 | *Bcl-2* | MBS701787 | MyBiosource | San Diego, CA | USA |
| 26 | *Survivin* | MBS030425 | MyBiosource | San Diego, CA | USA |
| 27 | *mTOR* | MBS2505637 | MyBiosource | San Diego, CA | USA |
| 28 | *PI3K* | MBS268899 | MyBiosource | San Diego, CA | USA |
| 29 | *Total protein* | MBS2540545 | MyBiosource | San Diego, CA | USA |
| 30 | *Albumin* | MBS730500 | MyBiosource | San Diego, CA | USA |
| 31 | *Globulin* | MBS2703040 | MyBiosource | San Diego, CA | USA |
| 32 | *Alanine transaminase* | MBS264333 | MyBiosource | San Diego, CA | USA |
| 33 | *Aspartate transaminase* | MBS450720 | MyBiosource | San Diego, CA | USA |
| **Cell line** | | **CAS No** | **Source** | **City** | **Country** |
| 34 | MCF-7 | ATCC HTB-22, RRID: CVCL_0031 | American Type Culture Collection | Manassas, VA | USA |
| 35 | MDA-MB-231 | ATCC CRM-HTB-26, RRID: CVCL_0062 | American Type Culture Collection | Manassas, VA | USA |
| 36 | SK-BR-3 | ATCC HTB-30, RRID: CVCL_0033 | American Type Culture Collection | Manassas, VA | USA |
| 37 | T47D | ATCC CRL-2865, RRID: CVCL_0553 | American Type Culture Collection | Manassas, VA | USA |

**Table ST2** Primer information for RT-PCR analysis

| **S.No** | **Primer name** | **Primer sequence** |
| --- | --- | --- |
| 1 | BCL2 | F: CAT GTG TGT GGA GAG CGT CAA |
|  |  | R: GCC GGT TCA GGT ACT CAG TCA |
| 2 | BAX | F: GTG GTT GCC CTC TTC TAC TTT GC |
|  |  | R: GAG GAC TCC AGC CAC AAA GAT G |
| 3 | P53 | F: GCC CCT CCT CAG CAT CTT AT |
|  |  | R: CTG TTC CGT CCC AGT AGA TT |
| 4 | CASP3 | F: CAT GGA AGC GAA TCA ATG GAC T |
|  |  | R: CTG TAC CAG ACC GAG ATG TCA |
| 5 | GAPDH | F: TGG GGA AGG TGA AGG TCG G |
|  |  | R: GGG ATC TCG CTC CTG GAA G |

| **Samples** | **Rct (Ω·cm^2^)** | **Remark** |
| --- | --- | --- |
| **ZnS** | ~6800 | Highest R_ct_; sluggish charge transfer due to poor conductivity. |
| **ZSS(0.25)** | ~1050 | Slightly less optimal than ZSS(0.20) and (0.15). |
| **ZSS(0.20)** | ~860 | Among the lowest; efficient charge separation and transport. |
| **ZSS(0.15)** | ~820 | Lowest R_ct_; best interfacial charge transport. |
| **ZSS(0.10)** | ~1150 | Higher R_ct_; performance starts to drop at low Zn doping. |
| **ZSS(0.05)** | ~1650 | Suboptimal composition; less favorable charge transfer. |
| **Ag_2_S** | ~5100 | High R_ct_ due to sluggish interfacial kinetics and higher resistance. |

**Table ST3** Charge transfer resistance (Rct) values were obtained from equivalent circuit fitting of the EIS data.

**Methods**

1. ***In-vitro studies***
   1.1. ***Cell culture***

The MCF-7 human breast adenocarcinoma cell line (ATCC HTB-22, RRID: CVCL_0031) was obtained from the American Type Culture Collection (ATCC, Manassas, VA, USA). Cells were maintained in Dulbecco’s Modified Eagle Medium (DMEM), supplemented with 10% fetal bovine serum (FBS) and 1% penicillin-streptomycin solution. Cultures were incubated at 37 °C in a humidified atmosphere containing 5% CO_2_. All cell handling procedures were carried out under aseptic conditions. Once the cultures reached approximately 80% confluence, cells were passaged. For experimental treatments, cells were seeded according to the requirements of each assay, as detailed in the respective experimental sections.

1.2. ***Fluorescent staining***

Following 24-hour treatment with ZSS nanostructures, both adherent and floating MCF-7 cells were carefully collected and washed with ice-cold 1× PBS to remove residual medium and treatment agents. The cells were subsequently fixed with 1 mL of 4% paraformaldehyde for 20 minutes at room temperature to maintain cellular morphology. After fixation, cells were rinsed again with PBS to eliminate any excess fixative. For assessment of nuclear integrity and oxidative stress, cells were stained with 4′,6-diamidino-2-phenylindole (DAPI) to detect nuclear condensation and fragmentation, and with dihydroethidium (DHE) to visualize intracellular ROS, following the manufacturer’s recommended protocols. Fluorescence imaging was performed using a Nikon Ti-E inverted fluorescence microscope equipped with appropriate excitation and emission filters for DAPI and DHE. High-resolution images were acquired at 10× magnification ^1^.

1.3. ***Gene expression analysis***

Total RNA was isolated from MCF-7 cells treated with ZSS nanostructures and cisplatin using the TRIzol reagent, following the manufacturer’s guidelines (Thermo Fisher Scientific, South Korea). For cDNA synthesis, 2 µg of RNA was used in a 40 µL reverse transcription reaction containing 1× RT buffer, 5 mM MgCl_2_, 10 mM dithiothreitol (DTT), 5 µM oligo(dT) primers (15-18 bases), 1 mM dNTP mix, 1 U/µL RNase inhibitor, and 10 U/µL M-MLV reverse transcriptase. The reaction was initiated by heating at 70 °C for 10 minutes, followed by rapid cooling at 4 °C for 2 minutes, and then incubation at 37 °C for 60 minutes to complete the reverse transcription ^2^. Quantification of gene expression was performed using SYBR Green-based quantitative real-time PCR (qRT-PCR) on a ViiA^TM^ 7 Real-Time PCR system (Applied Biosystems). The expression levels of apoptosis-related genes, including BCL2, BAX, P53, and CASP3 were analyzed using primers obtained from TaKaRa (South Korea) (**Table ST2**). Relative gene expression was normalized against an internal control gene (GAPDH) and analyzed using the ΔΔCt method.

2. ***In-vivo studies***
   2.1. ***Animal care and Ethical Compliance***

All experimental procedures involving animals were carried out in accordance with protocols approved by the Institutional Animal Care and Use Committee (IACUC) of Jeonbuk National University, Republic of Korea. For *in-vivo* evaluation of anticancer activity, 6 week-old female athymic nude BALB/c (nu/nu) mice (average body weight ~20 g) were obtained from Daehan BioLink (Eumsung, Korea). The mice were housed in a specific pathogen-free (SPF) facility under controlled experimental conditions, including a constant temperature of 23 ± 3 °C, regulated humidity, and a 12-hour light/dark cycle. Animals had free access to standard rodent feed and clean water throughout the duration of the experiment. All the *in-vivo* experiments were conducted in compliance with ARRIVE guidelines to ensure scientific rigor and ethical standards.

2.2. ***Tumor induction***

After a one-week acclimation period, mice were pre-implanted subcutaneously with 17β-estradiol pellets (0.72 mg (60 day release) Thermo Fisher Scientific, Cat: NC9032951) 72 hours prior to tumor inoculation to support the estrogen-dependent MCF-7 phenotype. MCF-7 human breast adenocarcinoma cells were used to establish subcutaneous xenografts in immunodeficient athymic Nude BALB/c mice. MCF-7 cell suspension (1 × 10^5^ cells) was suspended in a 1:1 mixture of cold PBS and Corning Matrigel® Matrix, High Concentration (HC)/LDEV-free (Cat: 354248) and injected subcutaneously into the dorsal flank of each mouse to establish localized xenografts. The estradiol supplementation-maintained ER⁺ activity, while the Matrigel depot stabilized the inoculum and enhanced local engraftment, ensuring reproducible and well-localized tumor formation. Tumor growth was systematically monitored at predefined intervals. Measurements of tumor length (A) and width (B) were taken using a digital vernier caliper. Tumor volume (V) was calculated using the standard ellipsoid formula: **V = A × B^2^ / 2**. This calculation allowed for consistent and accurate tracking of tumor progression throughout the experimental period ^3^.

2.3. ***Elisa- Apoptotic marker (Pro/Anti) - Protein quantification***

The tumor tissues excised from mice were homogenized in ice-cold phosphate buffered saline (PBS) using a tissue homogenizer to ensure complete cell lysis and release of intracellular proteins. The resulting homogenates were centrifuged at 2000 rpm for 15 minutes at 4 °C, and the clear supernatants were carefully collected for the subsequent quantification of apoptotic markers. Protein levels of both pro-apoptotic (Bax (Cat. No: MBS2513810), Caspase-3 (Cat. No: MBS260710), and p53 (Cat. No: MBS355295)) and anti-apoptotic ((Bcl-2 (Cat. No: MBS701787), Survivin (Cat. No: MBS030425), mTOR (Cat. No: MBS2505637), and PI3K (Cat.No: MBS268899)) proteins were determined using commercially available ELISA kits procured from MyBiosource (San Diego, CA, USA), following the manufacturer’s guidelines. Briefly, 40 μL of sample diluent was added to each well designated for the tumor samples, while 50 μL of standard solution was added to the wells reserved for standard curve generation. Subsequently, 100 μL of horseradish peroxidase (HRP)-conjugated reagent was added to all wells, and the plate was incubated at 37 °C for 60 minutes to facilitate antigen-antibody binding. After incubation, the wells were washed five times with the provided wash buffer to remove unbound reagents. Thereafter, 50 μL each of Chromogen Solution A and Chromogen Solution B were added to every well, gently mixed, and incubated in the dark at 37 °C for 15 minutes to develop the color. The enzymatic reaction was terminated by the addition of 50 μL of stop solution, leading to a color change from blue to yellow. Absorbance was measured at 450 nm using a microplate reader, and the concentration of each apoptotic marker was calculated based on the standard curve derived from known reference concentrations ^4, 5^.

2.4. ***Liver functionality test***

To assess liver function, the excised liver tissues were first homogenized in chilled PBS using a tissue homogenizer to ensure uniform sample processing. The resulting homogenates were centrifuged at 2000 rpm for 15 minutes at 4 °C to separate the supernatant, which was then collected for subsequent biochemical evaluation. Key hepatic function indicators, including total protein (Cat. No: MBS2540545), albumin (Cat. No: MBS730500), globulin (Cat. No: MBS2703040), alanine transaminase (ALT, Cat. No: MBS264333), and aspartate transaminase (AST, Cat. No: MBS450720), were measured using commercially available colorimetric assay kits (MyBiosource, San Diego, CA, USA). All analyses were conducted in strict accordance with the manufacturers’ protocols to maintain consistency and analytical accuracy ^4, 6^.

**Figure SF1** Particle size distribution of ZSS(0.15)

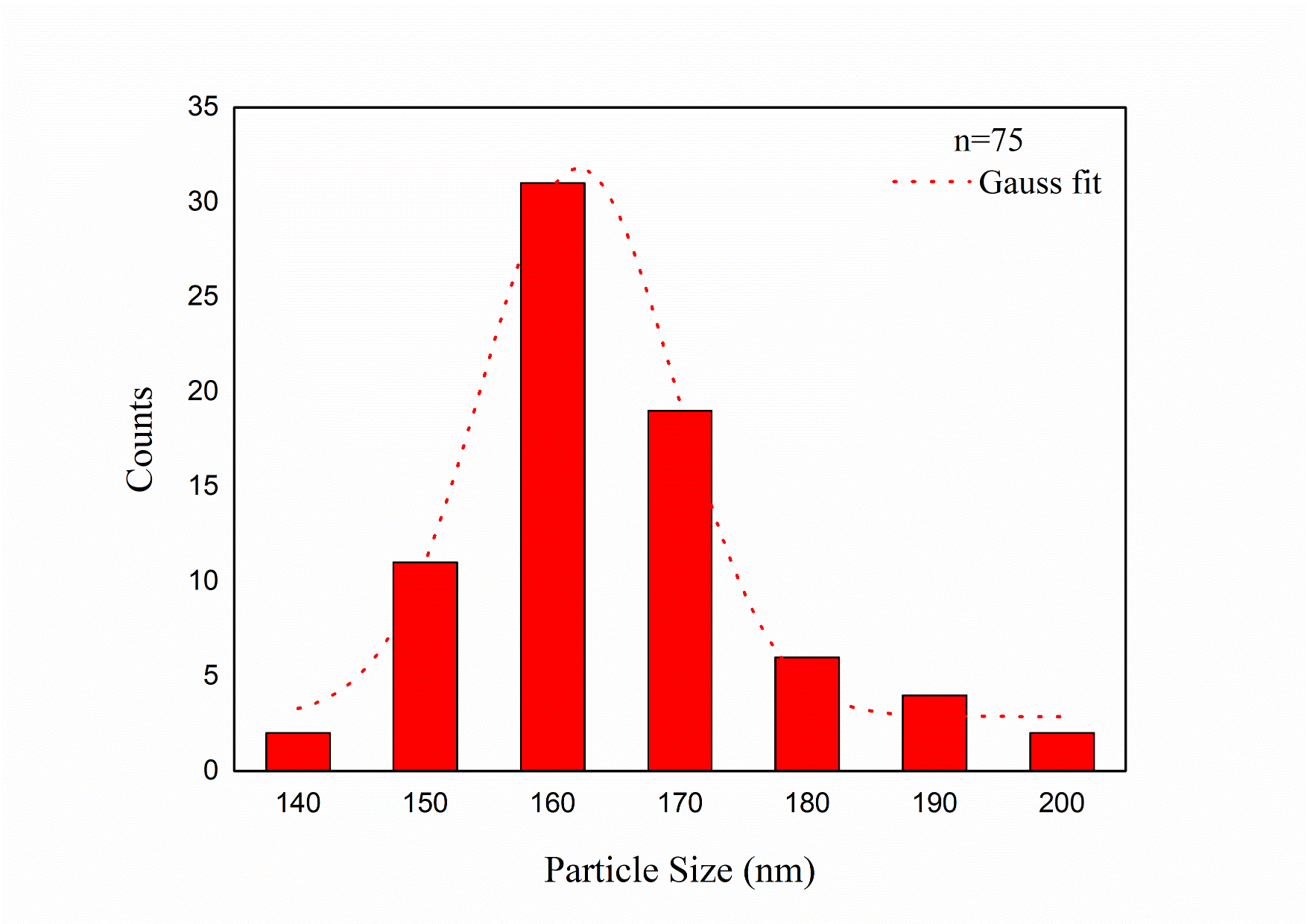

**Figure SF2** Photothermal conversion plot

**Photothermal Conversion Efficiency (η) Formula**: $\boldsymbol{\eta}\mathbf{=}\frac{\boldsymbol{hS}\left( \boldsymbol{T}_{\boldsymbol{max}}\boldsymbol{-}\boldsymbol{T}_{\boldsymbol{amb}} \right)\boldsymbol{-}\boldsymbol{Q}_{\boldsymbol{dis}}}{\boldsymbol{I(1}\boldsymbol{-}\boldsymbol{10}^{\boldsymbol{-}\boldsymbol{A660}}\boldsymbol{)}}$

**Where:**

| ***h -*** Heat transfer coefficient  ***S-*** Surface area of the container (m^2^) | | To determine hS,  $\theta= \frac{T\left( t \right)-T_{s}}{T_{max}- T_{S}}$;  Slope of ln(θ)=−hS/mC [m= mass of water (0.0002 kg; C= heat capacity 4184 J/kg K]; |
| --- | --- | --- |
| **T_max_** | Maximum temperature reached by the sample (K) | 56 °C (329.15 K) |
| **T_amb_** | Ambient temperature (K) | 25 °C (298.15 K) |
| **Q_dis_** | Heat dissipated from light absorbed by solvent/container | negligible |
| **I** | Laser power density | 0.5 (W/cm^2^) |
| **A660** | Absorbance of sample at 660 nm | 0.344 |

*Calculate θ and ln(θ)*

$$\theta= \frac{T\left( t \right)-T_{s}}{T_{max}- T_{S}}$$

where,
T(t) = temperature at time t,
T_s_ = ambient temperature,

T_max_ = maximum temperature reached.

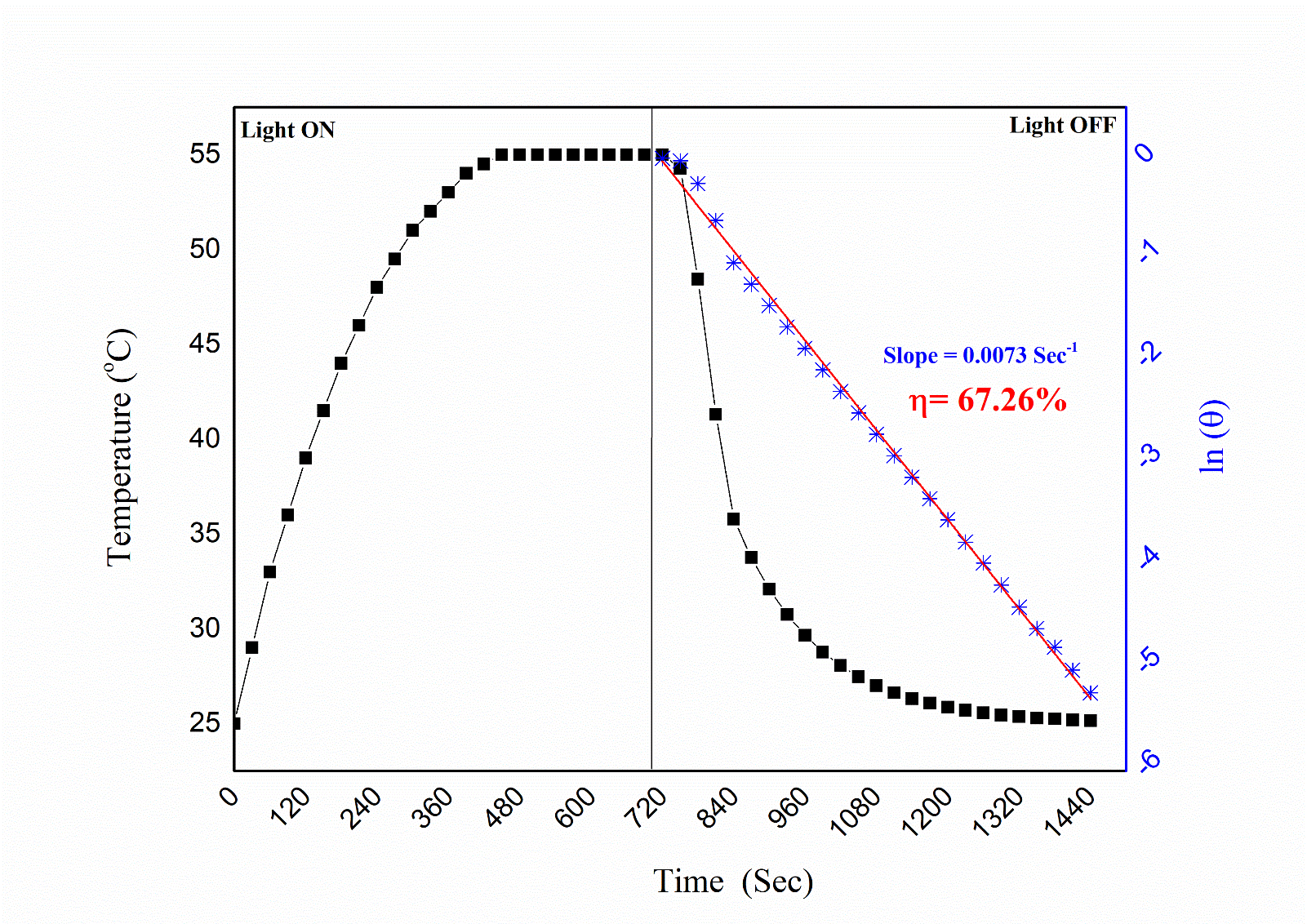

***Serum stability evaluation***

The colloidal stability of ZSS(0.15) under physiologically relevant conditions was evaluated by monitoring changes in hydrodynamic size, polydispersity index (PDI), and ζ-potential over time in PBS (pH 7.4) and PBS supplemented with 10% fetal bovine serum (FBS) (**Figure SF3**). In both media, ZSS(0.15) exhibited a gradual and moderate increase in hydrodynamic diameter over 24 h, while maintaining PDI values below 0.35, indicating the absence of severe aggregation. The presence of serum proteins resulted in a slightly larger size increase, consistent with the formation of a protein corona, but without compromising colloidal stability. Additionally, the ζ-potential remained consistently negative with only minor variation over time in both PBS and PBS + 10% FBS, suggesting preservation of surface charge characteristics. Collectively, these results demonstrate that ZSS(0.15) maintains acceptable colloidal stability in serum-containing environments, supporting its suitability for biological and *in-vivo* applications.

**Figure SF3** **(a)** Time-dependent changes in hydrodynamic size measured by dynamic light scattering (DLS) for ZSS(0.15) dispersed in PBS (pH 7.4) and PBS containing 10% fetal bovine serum (FBS); **(b)** Corresponding polydispersity index (PDI) and **(c)** ζ-potential of ZSS(0.15) measured at 0 h and 24 h in PBS and PBS + 10% FBS

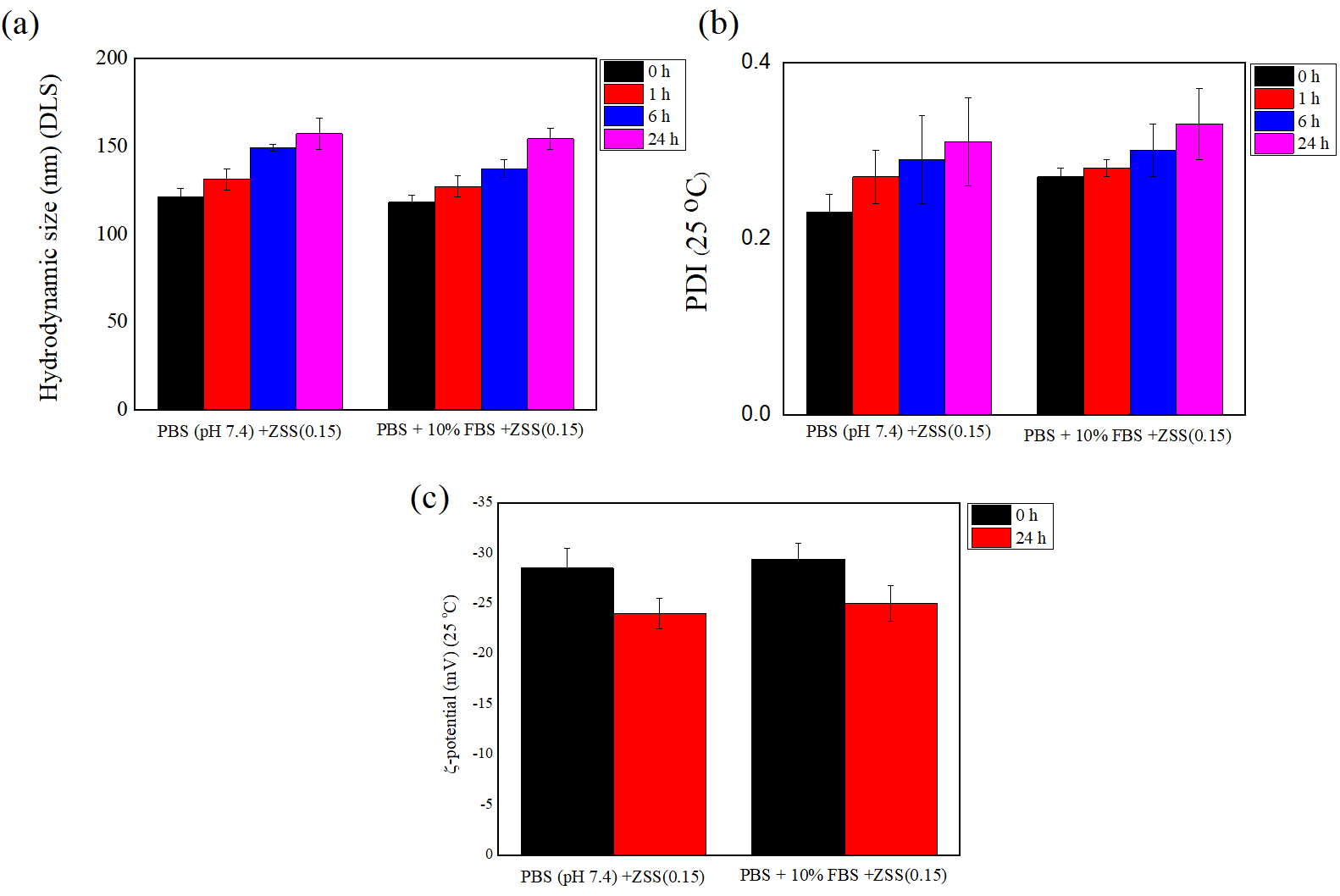

**REFERENCE**

(1) Mohan, H.; Ramalingam, V.; Lim, J.-M.; Lee, S.-W.; Kim, J.; Lee, J.-H.; Park, Y.-J.; Seralathan, K.-K.; Oh, B.-T. E-waste based graphene oxide/V2O5/Pt ternary composite: Enhanced visible light driven photocatalyst for anti-microbial and anti-cancer activity. *Colloids and Surfaces A: Physicochemical and Engineering Aspects* **2020**, *607*, 125469. DOI: <https://doi.org/10.1016/j.colsurfa.2020.125469>.

(2) Varunkumar, K.; Anusha, C.; Saranya, T.; Ramalingam, V.; Raja, S.; Ravikumar, V. Avicennia marina engineered nanoparticles induce apoptosis in adenocarcinoma lung cancer cell line through p53 mediated signaling pathways. *Process Biochemistry* **2020**, *94*, 349-358.

(3) Liu, G.; Gao, N.; Zhou, Y.; Nie, J.; Cheng, W.; Luo, M.; Mei, L.; Zeng, X.; Deng, W. Polydopamine-Based “Four-in-One” Versatile Nanoplatforms for Targeted Dual Chemo and Photothermal Synergistic Cancer Therapy. In *Pharmaceutics*, 2019; Vol. 11.

(4) Rizzk, Y. W.; El-Deen, I. M.; Behery, M. E.; Mourad, A. A. E.; Mohammed, F. Z. In vivo anticancer study of sodium 2-[(4-oxidobenzylidene)amino]-6H-1,3,4-thiadiazine-5-olate against Ehrlich ascites carcinoma via targeting PI3K/mTOR pathway. *Chemical Biology & Drug Design* **2024**, *103* (2), e14452. DOI: <https://doi.org/10.1111/cbdd.14452> (acccessed 2025/04/27).

(5) Younis, N. S.; Ghanim, A. M. H. The Protective Role of Celastrol in Renal Ischemia-Reperfusion Injury by Activating Nrf2/HO-1, PI3K/AKT Signaling Pathways, Modulating NF-κb Signaling Pathways, and Inhibiting ERK Phosphorylation. *Cell Biochemistry and Biophysics* **2022**, *80* (1), 191-202. DOI: 10.1007/s12013-022-01064-6.

(6) Bergmeyer, H.-U. *Methods of enzymatic analysis*; Elsevier, 2012.
